# Testing the CRISPR-Cas9 and *glmS* ribozyme systems in *Leishmania tarentolae*

**DOI:** 10.1101/2020.07.23.218073

**Authors:** Gino L. Turra, Luzia Schneider, Linda Liedgens, Marcel Deponte

## Abstract

*Leishmania* parasites include important pathogens and model organisms and are even used for the production of recombinant proteins. However, functional genomics and the characterization of essential genes are often limited in *Leishmania* because of low-throughput technologies for gene disruption or tagging and the absence of components for RNA interference. Here, we tested the T7 RNA polymerase-dependent CRISPR-Cas9 system by Beneke *et al*. and the *glmS* ribozyme-based knock-down system in the model parasite *Leishmania tarentolae*. We successfully deleted two reference genes encoding the flagellar motility factor Pf16 and the salvage-pathway enzyme adenine phosphoribosyltransferase, resulting in immotile and drug-resistant parasites, respectively. In contrast, we were unable to disrupt the gene encoding the mitochondrial flavoprotein Erv. Cultivation of *L. tarentolae* in standard BHI medium resulted in a constitutive down-regulation of an episomal *mCherry-glmS* reporter by 40 to 60%. For inducible knock-downs, we evaluated the growth of *L. tarentolae* in alternative media and identified supplemented MEM, IMDM and McCoy’s 5A medium as candidates. Cultivation in supplemented MEM allowed an inducible, glucosamine concentration-dependent down-regulation of the episomal *mCherry-glmS* reporter by more than 70%. However, chromosomal *glmS*-tagging of the genes encoding Pf16, adenine phosphoribosyltransferase or Erv did not reveal a knock-down phenotype. Our data demonstrate the suitability of the CRISPR-Cas9 system for the disruption and tagging of genes in *L. tarentolae* as well as the limitations of the *glmS* system, which was restricted to moderate efficiencies for episomal knock-downs and caused no detectable phenotype for chromosomal knock-downs.

## Introduction

Kinetoplastid parasites cause important diseases, such as sleeping sickness, Chagas disease as well as cutaneous, mucocutaneous or visceral leishmaniasis^1,2^. Furthermore, kinetoplastid parasites are valuable unicellular model organisms and have led to seminal discoveries in molecular cell biology, including RNA editing^3^ or the structure of the glycosylphosphatidylinositol anchor^4^. The gecko parasite *Leishmania tarentolae*, for example, is nonpathogenic for humans, grows fast, serves as a valuable model system for RNA editing^5^ and mitochondrial protein import^6^ and is also used for the constitutive or tetracycline-induced production of recombinant proteins^7,8^. However, gene disruption or tagging in diploid *L. tarentolae* so far relied on rather laborious traditional genetics. The recent introduction of the CRISPR-Cas9 technology in kinetoplastid parasite research^9–11^ now provides the opportunity to overcome this limitation, especially taking into account that a high-throughput method for gene disruption or tagging was successfully established in *L. major, L. mexicana* and *L. donovani*^12–14^.

Molecular tools to alter intracellular RNA or protein concentrations are crucial for functional genomics, in particular, to address essential genes that cannot be deleted. One common approach is to down-regulate a protein of interest with the help of a genetically encoded protein tag that is degraded in the absence of a stabilizing ligand^15^. However, protein tagging may affect the localization and function of the protein of interest and is sometimes even used to interfere with protein trafficking^16^. Furthermore, stabilization of the protein tag usually requires the long-term addition of an expensive ligand to the cell culture medium. An alternative approach is to down-regulate the protein-encoding RNA of interest by RNA interference (RNAi)^17,18^. RNAi is a widely used method for post-transcriptional gene repression in eukaryotes but requires a specific RNAi machinery for RNA degradation^19^. This machinery has been lost in many eukaryotes, e.g., in several important apicomplexan and kinetoplastid parasites^20–22^. A transcriptional knock-down approach is to generate transgenic cells with a T7 RNA polymerase and a tetracycline-controlled *trans*-activator, which is a fusion protein between a tetracycline repressor and a transactivation domain^23^. This system also requires an artificial operon that comprises a T7 promoter, the gene of interest and a tetracycline operator. Depending on the type of transactivation domain, the *trans*-activator binds to the operator and activates the T7 RNA polymerase either in the absence or presence of tetracycline^24^. One disadvantage of this versatile transcriptional system is that tetracycline or related antibiotics may also block translation in mitochondria or plastids in a time- and concentration-dependent manner^25^. A fourth knock-down approach utilizes genetically encoded fusion constructs between an RNA of interest and the *glmS* ribozyme, which is derived from a part of the 5’ untranslated region of the *Bacillus subtilis* glucosamine-6-phosphate synthase gene^26^. The *glmS* ribozyme is activated by glucosamine-6-phosphate or, to a lesser degree, by the precursor glucosamine, and is inhibited by glucose-6-phosphate^26,27^. Hence, fusion constructs between the RNA of interest and the *glmS* ribozyme are destabilized by the addition of glucosamine. The amino group of bound glucosamine-6-phosphate is part of the active site and participates in the self-cleavage of an AG-motif at the 5’-end of the ribozyme^26,28,29^. Replacement of the AG-motif with a CC-motif in the so-called *glmS* M9 mutant prevents self-cleavage and can be used as a negative control^26^. The advantages of the *glmS* ribozyme system are its simplicity and the low costs for glucosamine. However, depending on the investigated organism or cell system, high concentrations of glucosamine might be toxic^30^, and the activity of the *glmS* ribozyme might depend on the medium composition and the ratio between ribozyme activators and inhibitors^31^.

Here, we successfully adopted the LeishGEdit CRISPR-Cas9 method by Beneke et al.^12,32^ to generate knock-out and knock-in strains and to test the *glmS* ribozyme-based knock-down system in *L. tarentolae*. Deletion of the flagellar motility factor *PF16* resulted in immotile parasites, deletion of adenine phosphoribosyltransferase (*APRT*) rendered parasites resistant to 4-aminopyrazolopyrimidine, and deletion of the mitochondrial flavoprotein *ERV* was lethal. Using a non-invasive plate reader assay for monitoring fluorescent reporter proteins, we found that the medium composition influences the *glmS* ribozyme-mediated knock-down effect. Standard brain-heart-infusion (BHI) medium resulted in a constitutive down-regulation of an episomal *mCherry-glmS* reporter, whereas supplemented minimal essential medium (MEM) allowed a glucosamine concentration-dependent down-regulation. However, chromosomal *glmS*-tagging of the genes encoding Pf16, APRT or Erv did not result in a detectable knock-down phenotype.

## Results

### Generation of *L. tarentolae* knock-out strains using the CRISPR-Cas9 system

*L. tarentolae* promastigotes were transfected with plasmid pTB007 encoding T7 RNA polymerase and FLAG-tagged Cas9 as described previously for *L. major* and *L. mexicana*^12,13^ (Fig. 1a). Western blot analysis confirmed the presence of Cas9 (Fig. 1b) and the functionality of the T7 RNA polymerase was shown by fluorescence microscopy using an mCherry reporter under control of a T7 promoter (Fig. 1c). Furthermore, growth analyses revealed no detrimental effect for plasmid pTB007 in *L. tarentolae* (Fig. 1d).

**Figure 1.**
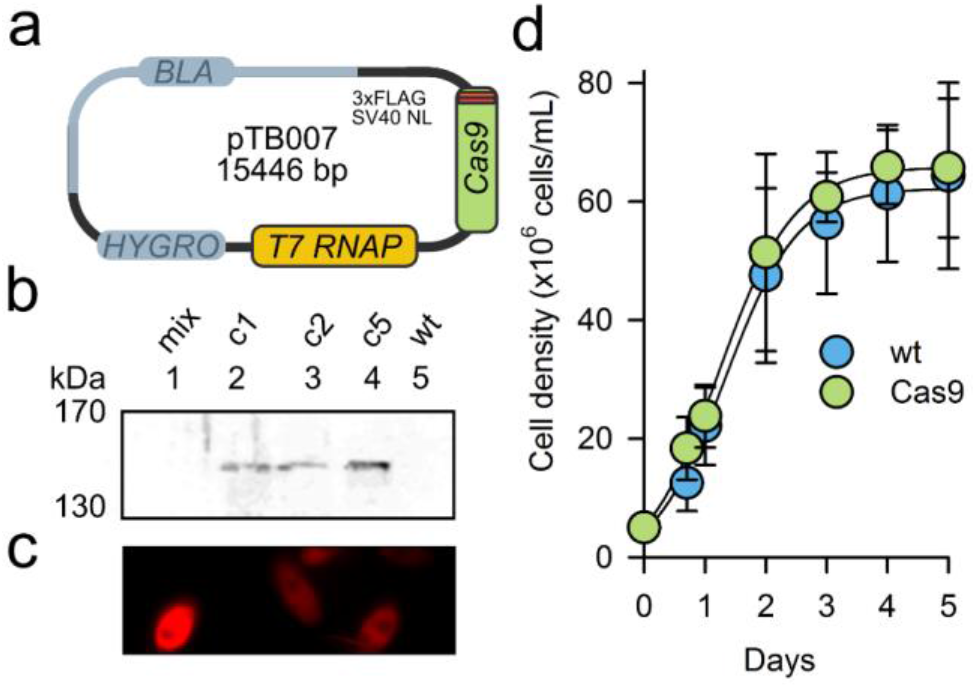
Validation of Cas9 and T7 RNA polymerase expression in *L. tarentolae*. (**a**) *L. tarentolae* promastigotes were transfected with plasmid pTB007 encoding T7 RNA polymerase (*T7 RNAP*) and triple FLAG-tagged Cas9 from *Streptococcus pyogenes* with a nuclear localization sequence (NL). Selection markers against β-lactam antibiotics (*BLA*) and hygromycin B (*HYGRO*) are indicated. (**b**) Western blot analysis with an anti-FLAG antibody confirmed the presence of tagged Cas9 in accordance with the calculated molecular mass of 160 kDa. Three clonal cell lines were obtained after selection with 100 μg/mL hygromycin. Wild-type cells (wt) served as a control. For each strain 5×10^6^ cells were boiled in Laemmli buffer and loaded per lane. (**c**) The functionality of the T7 RNA polymerase was tested using plasmid pLEXSY_IE-blecherry4 that encodes mCherry under the control of a T7 promoter. Fluorescence microscopy confirmed the presence of mCherry throughout the cytosol. (**d**) Growth curve analyses in BHI medium revealed no differences between the wild-type strain (wt) and *L. tarentolae* promastigotes carrying plasmid pTB007 (Cas9), which served as the parental strain for subsequent experiments. Data are the mean ± standard deviation of three biological replicates.

As proof-of-principle experiments and because of the expected characteristic phenotypes that could also serve as references for subsequent knock-down studies, we then deleted either the gene for the flagellar motility factor *PF16* or the adenine phosphoribosyltransferase (*APRT*) (Fig. 2). These modifications should render *L. tarentolae* promastigotes either immotile, due to the loss of *PF16*, or resistant to the pro-drug 4-aminopyrazolopyrimidine, due to the loss of *APRT*, as reported previously for *L. mexicana*^12^ and *L. donovani*^33,34^, respectively. Briefly, *L. tarentolae* parasites with T7 RNA polymerase and Cas9 were used to drive the *in vivo* transcription of two different gene-specific single guide RNAs (sgRNA) for the excision of the gene of interest^12,32^. The two sgRNA-encoding DNAs were generated by PCR using an universal sgRNA antisense primer and a sense primer that encodes the 24 nucleotide T7 promoter, the site-specific 20 nucleotide guide sequence and a 20 nucleotide sequence that is complementary to the universal sgRNA antisense primer. Plasmid pTPuro was used to amplify a puromycin resistance cassette by PCR using primers with 30 nucleotide homology regions upstream and downstream of the first and second sgRNA-induced double-strand break, respectively^12,32^. Parasites with plasmid pTB007 were co-transfected with the three PCR products (encoding the two sgRNAs for the excision and the puromycin cassette for the repair of the double-strand breaks) and selected on BHI agar plates with 20 μg/mL puromycin. Colonies appeared after 7-14 days and were transferred to liquid culture containing 20 μg/mL puromycin. PCR and motility analyses confirmed the loss of *PF16* after a single round of selection, resulting in dividing but completely immotile promastigotes (Fig. 2b,c). PCR analysis and dose-response curves for 4-aminopyrazolopyrimidine also confirmed the loss of *APRT* after a single round of selection, resulting in drug-resistant parasites (Fig. 2e,f).

**Figure 2.**
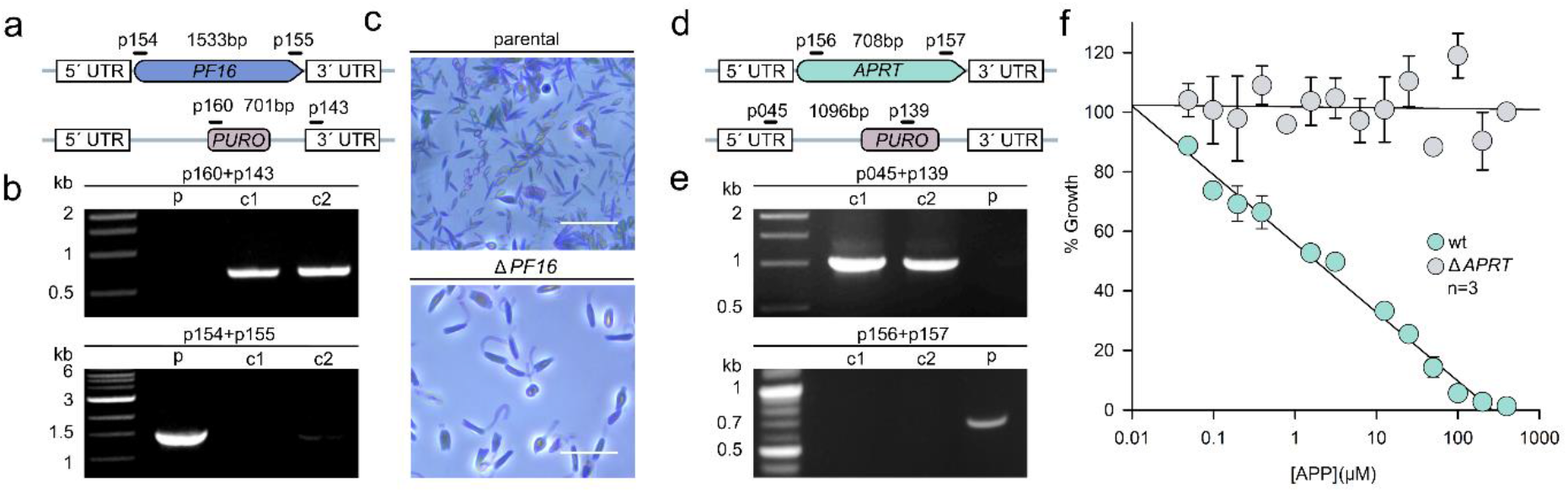
Generation and validation of *L. tarentolae* knock-out parasites using the CRISPR-Cas9 system. (**a**) Schematic representation of the replacement of the chromosomal *PF16* locus with a cassette that encodes the puromycin *N*-acetyltransferase (*PURO*). Only one of both *L. tarentolae* gene copies is shown. Primer positions and expected product sizes from analytical PCR reactions are highlighted. (**b**) PCR analyses of genomic DNA of the parental strain (p) and of two clonal homozygous knock-out cell lines after transfection and selection. PCR products were analyzed by agarose gel electrophoresis confirming the expected product sizes from panel a. Primer pairs are indicated on top of each gel. (**c**) *L. tarentolae* motility as observed by phase contrast microscopy. Twenty-five consecutive frames recorded within 23.5 seconds were stacked in one single image. Motile parental cells with typical movement are shown on top and immotile homozygous *pf16* knock-out cells (Δ*PF16*) are shown below. Scale bars: 10 μm. (**d**) Schematic representation of the replacement of the chromosomal *APRT* locus. Only one of both gene copies is shown. Primer positions and expected product sizes from analytical PCR reactions are highlighted. (**e**) PCR analyses confirmed the replacement of *APRT* for two clonal homozygous knock-out cell lines. (**f**) Dose-response curves for 4-aminopyrazolopyrimidine (APP) revealed drug resistance for homozygous *aprt* knock-out parasites (Δ*APRT*) in contrast to susceptible wild-type parasites (wt).

We previously showed that the gene encoding the mitochondrial flavoenzyme Erv cannot be deleted in the related human pathogen *L. infantum* using standard genetics^35^. This enzyme is thought to play an important role in mitochondrial protein import in kinetoplastid parasites^6,36–38^. We therefore wanted to test whether the CRISPR-Cas9 method might allow us to select knock-out parasites that can be missed in standard genetic experiments (for example, slow-growing parasites that are usually overgrown by plasmid-containing wild-type parasites without negative selection). Using the same strategy as for *PF16* and *APRT*, we were unable to delete *ERV* in *L. tarentolae*. No colonies appeared following transfection with the correct PCR products and selection with puromycin in two independent experiments. Hence, *ERV* is most likely essential in *Leishmania* parasites. In summary, we showed that the PCR- and T7 RNA polymerase-based CRISPR-Cas9 system by Beneke et al. can be also used in *L. tarentolae*. Furthermore, in accordance with previous findings in other *Leishmania* species, we show that (i) *PF16* is crucial for promastigote motility, (ii) deletion of *APRT* confers resistance towards the pro-drug 4-aminopyrazolopyrimidine and (iii) *ERV* is refractory to disruption.

### Limitations of initial experiments with *GFP-glmS* and BHI medium

In order to test the *glmS* ribozyme system in *L. tarentolae*, we initially chose to quantify the potential down-regulation of plasmid-encoded green fluorescent protein (GFP) in intact cells. A *GFP-glmS* fusion construct was cloned into vector pX, and transfected clonal *L. tarentolae* cell lines were subsequently cultured in standard BHI medium with or without 10 mM glucosamine (Fig. S1a). A rather weak fluorescence was observed for control cultures without glucosamine (Fig. S1b). Although parasite cultures with glucosamine appeared to have an even 50% lower fluorescence, we could not obtain statistically meaningful quantitative data from three biological replicates because of the variable fluorescence of the control culture. The rather weak GFP fluorescence in the absence of external glucosamine might have resulted from an unfavorable, medium-dependent ratio between *glmS* activators and inhibitors. The residual fluorescence in the presence of glucosamine might have been caused by an inefficient down-regulation or by the mitochondrial autofluorescence (which was previously shown to overlap with the fluorescence of GFP and to give a false positive signal, particularly at a low signal-to-noise ratio^39^). In order to overcome these potential limitations, we tested alternative media and used mCherry as a fluorescent reporter in subsequent experiments.

### Identification of alternative cell culture media

To address a putative effect of the medium composition on the *glmS* ribozyme system, as reported in yeast^31^, we first had to establish an alternative cell culture protocol. Therefore, we analyzed the growth of *L. tarentolae* in six different (defined) liquid media that are also available as SILAC media for potential future knock-down studies in combination with quantitative mass spectrometry. The media were supplemented with different concentrations of folic acid, hemin and fetal bovine serum (FBS) resulting in 216 different media as described in the Materials and Methods section. DMEM, DMEM/F12 and RPMI-1640 media as well as media without FBS were altogether excluded from further analysis because parasites died within one to three days regardless of the supplement concentrations. In contrast, parasites from selected FBS-containing cultures with supplemented MEM, IMDM and McCoy’s 5A medium survived. These media were subsequently analyzed in more detail and compared to standard BHI medium as a control. In general, parasite growth was better with 10% FBS compared to 5% FBS. Highest maximum cell densities were observed in McCoy’s 5A and BHI medium followed by IMDM and MEM (Fig. 3a). The morphology and motility of parasites in FBS-containing MEM, IMDM and McCoy’s 5A medium were similar to the control in BHI medium (Fig. 3b). Attempts to replace FBS with 0.5% AlbuMax II resulted in cell aggregation and replacement with dialyzed FBS lead to cell death. Hence, we could not identify a suitable medium with dialyzed FBS for potential SILAC experiments. Nevertheless, we decided to test the *glmS* ribozyme system in supplemented MEM containing 5% FBS, because MEM contains 50% less glucose (the precursor of the *glmS* activator glucosamine and the inhibitor glucose-6-phosphate) than BHI medium and has the lowest glucose concentration among the three defined media (1.0, 3.0 and 4.5 g/L in MEM, McCoy’s 5A medium and IMDM, respectively). The low toxicity of glucosamine in MEM was another positive factor for subsequent *glmS* studies. Single passages of parasite cultures to MEM containing up to 15 mM glucosamine had no toxic effect and concentrations above 15 mM glucosamine reduced the final cell density by only one-fourth (Fig. 3c). In summary, we identified supplemented MEM, IMDM and McCoy’s 5A medium with 5 or 10% FBS as suitable liquid media for the growth of *L. tarentolae* and selected MEM supplemented with 10 μg/mL hemin chloride, 25 mM HEPES and 5% FBS for subsequent *glmS* studies because of its low glucose content and glucosamine toxicity profile.

**Figure 3.**
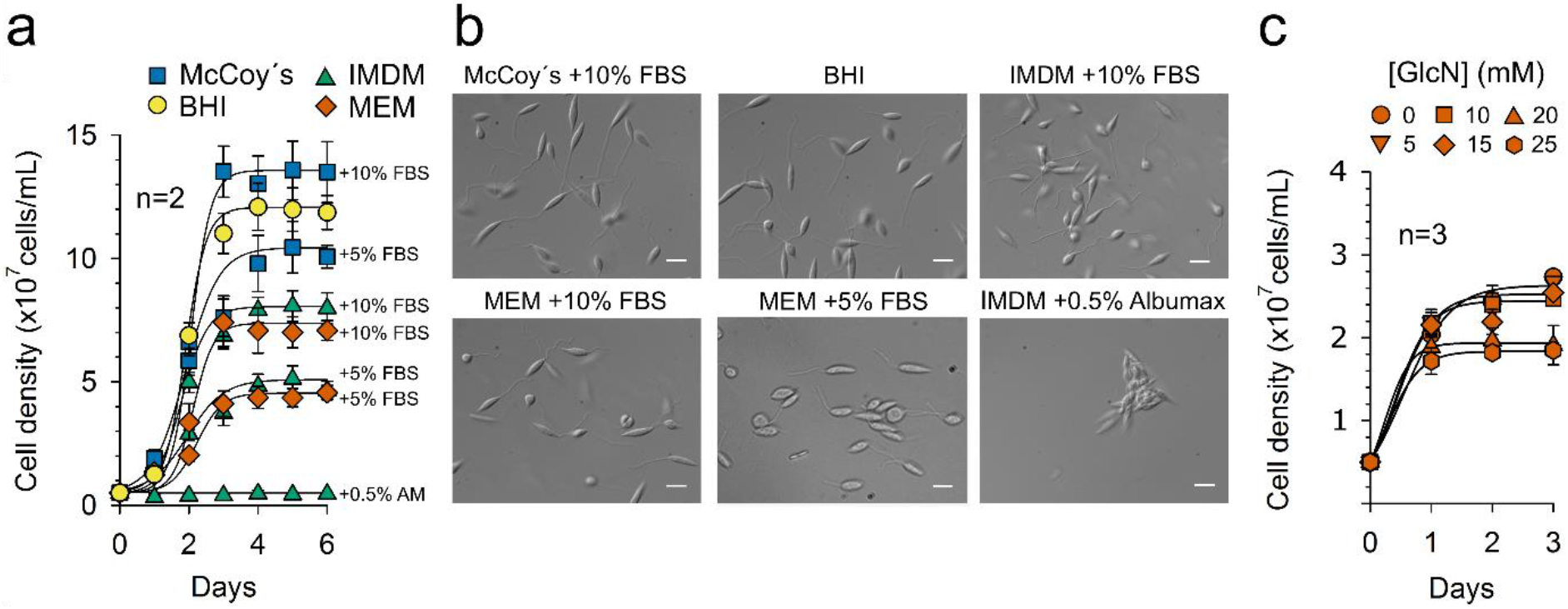
Growth rates of *L. tarentolae* in different liquid media. (**a**) Growth curve analysis for *L. tarentolae* liquid cultures using the indicated alternative media. All media contained 10 μg/mL hemin chloride. MEM and McCoy’s were also supplemented with 25 mM HEPES. Data points represent the mean ± standard deviation from two independent biological replicates. Sigmoidal fits were generated in SigmaPlot 13. FBS, fetal bovine serum; AM, AlbuMax II. (**b**) Representative microscopy images of *L. tarentolae* promastigotes from panel a that were grown in the indicated media. Images were taken during the mid-logarithmic phase. Scale bars: 10 μm. (**c**) Growth curve analysis for *L. tarentolae* liquid cultures in MEM that was supplemented with 10 μg/mL hemin chloride, 25 mM HEPES, 5% FBS and the indicated concentrations of glucosamine (GlcN). Data points represent the mean ± standard deviation from three independent biological replicates. Sigmoidal fits were generated in SigmaPlot 13.

### An inducible episomal *glmS* ribozyme knock-down system in *L. tarentolae*

To overcome the potential limitations described above, we cloned an *mCherry-glmS* fusion construct in vector pX and analyzed the *glmS* ribozyme system for transfected clonal cell lines of *L. tarentolae* that were cultured in supplemented MEM (Fig. 4a). As a control, we also fused *mCherry* to the inactive *glmS* M9 mutant to study the potential effect of endogenous *glmS* activators and inhibitors. The mCherry fluorescence was quantified in a microplate reader after three passages in supplemented MEM with or without 5 or 10 mM glucosamine (Fig. 4b). This protocol was chosen to dilute the intracellular signal from stable mCherry that was already synthesized before the addition of glucosamine. In contrast to single glucosamine treatments in Fig. 3c, the repeated addition of 10 mM glucosamine resulted in a time-dependent growth retardation (Fig. 4c), which was compensated in the fluorescence measurements by seeding the same number of parasites per well. The wild-type *glmS* construct was reproducibly down-regulated in the presence of glucosamine in a concentration-dependent manner (Fig. 4d). Furthermore, even in the absence of external glucosamine, the mCherry signal of the culture with the wild-type *glmS* construct was about 35% lower than the fluorescence of the *glmS* M9 control, suggesting that endogenous glucosamine-6-phosphate causes an intrinsic *glmS* ribozyme activation and down-regulation of mCherry. The addition of 10 mM glucosamine to the medium significantly increased the difference between the fluorescence of cultures with wild-type *glmS* and the *glmS* M9 control and resulted in a down-regulation of mCherry by more than 70%. In summary, we established an inducible *glmS* knock-down system for a plasmid-encoded reporter in *L. tarentolae* using supplemented MEM as an alternative cell culture medium.

**Figure 4.**
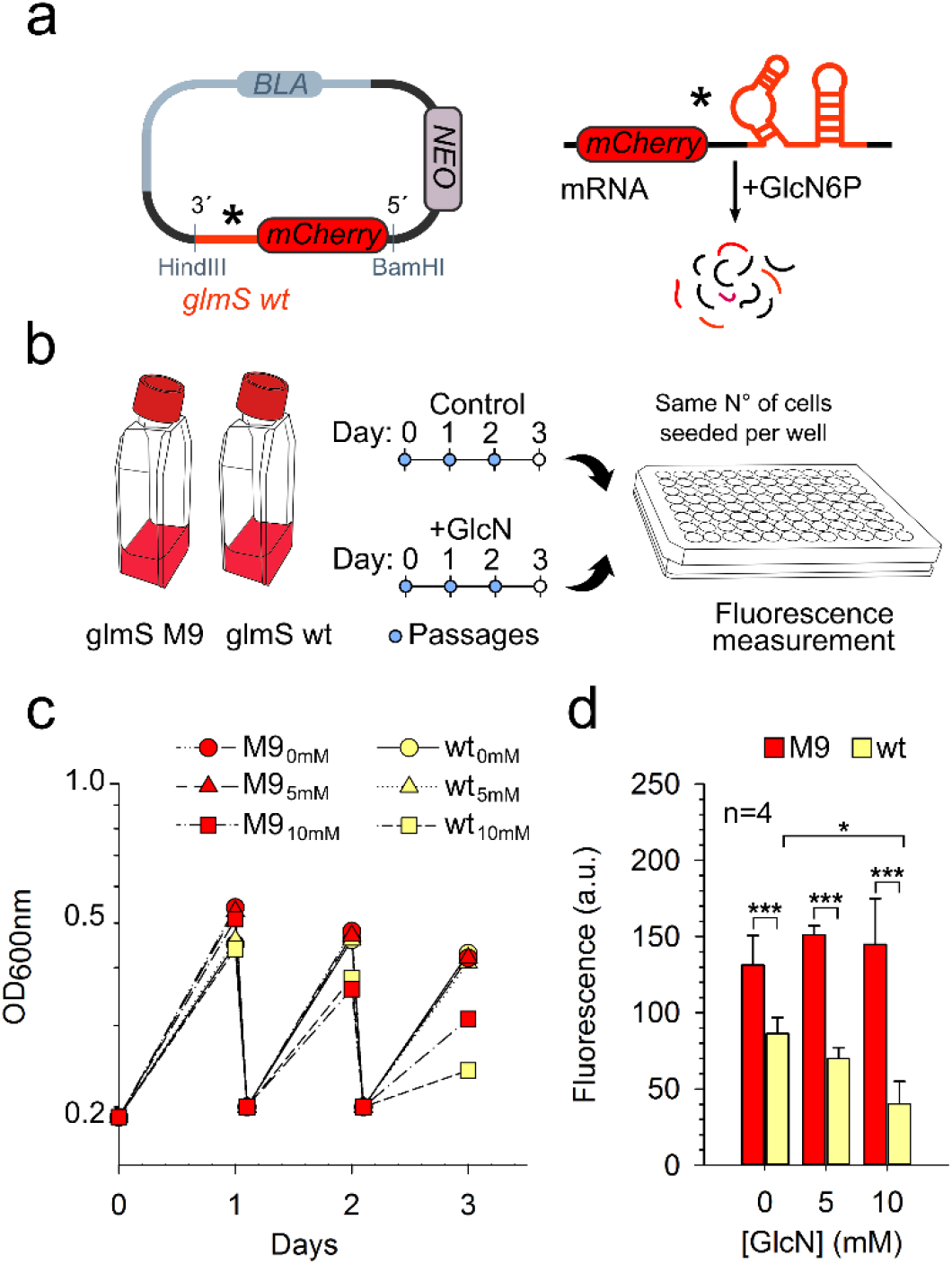
Inducible *mCherry-glmS* ribozyme knock-down in *L. tarentolae* that was grown in MEM. (**a**) Schematic overview of the used *mCherry-glmS* ribozyme construct in vector pX. The encoded mRNA should be cleaved and subsequently degraded in the presence of the activating ligand glucosamine-6-phosphate (GlcN6P). The asterisk indicates that an inactive *glmS* M9 mutant can be used as a negative control. (**b**) Schematic overview of the knock-down cell culture protocol. Clonal cell lines with wild-type (wt) or mutant (M9) *mCherry-glmS* ribozyme were grown for three days with or without 10 mM glucosamine (GlcN). Parasites were diluted every day in fresh medium to an initial OD of 0.2 before the mCherry fluorescence was analyzed in intact cells at day three. (**c**) Representative growth curve analysis for *L. tarentolae* liquid cultures with wt or M9 *mCherry-glmS* ribozyme according to the scheme in panel b. All cell lines were grown in parallel. MEM was supplemented with 10 μg/mL hemin chloride, 25 mM HEPES, pH 7.4, 5% FBS and the indicated concentration of glucosamine (0, 5 or 10 mM). (**d**) Normalized mCherry fluorescence of cell cultures with 0, 5 or 10 mM glucosamine (GlcN) from panel c at day three. All data points are the mean ± standard deviation from four biological replicates. Statistical analyses were performed in SigmaPlot 13 using the one-way ANOVA method (* p < 0.05; *** p < 0.001).

### A constitutive episomal *glmS* ribozyme knock-down system in *L. tarentolae*

To address whether the *glmS* ribozyme system can be also used in BHI medium, we repeated the experiments for plasmid-encoded wild-type and mutant *mCherry-glmS* according to the protocol in Fig. 4b using standard BHI medium instead of supplemented MEM. The mCherry fluorescence of cultures with the wild-type *glmS* construct was lowered by up to two-thirds compared to the *glmS* M9 control (Fig. 5). The results were highly reproducible for two independent clonal cell lines and among biological replicates. External glucosamine seemed to have no effect on the mCherry fluorescence, suggesting that internal *glmS* activator(s) and inhibitor(s) overlay the impact of external glucosamine in BHI medium. A constitutive intrinsic down-regulation of wild-type *glmS* constructs in BHI medium without external glucosamine also explains (i) a passage-independent constant mCherry fluorescence at days 1, 2 and 3 (data not shown) and (ii) the rather low GFP fluorescence in our initial experiments that were performed without a *glmS* M9 control (Fig. S1). In summary, the *glmS* ribozyme system can be used in BHI medium for constitutive knock-down experiments using plasmid-encoding genes. Control experiments with *glmS* M9 constructs enable a quantification of the degree of constitutive down-regulation.

**Figure 5.**
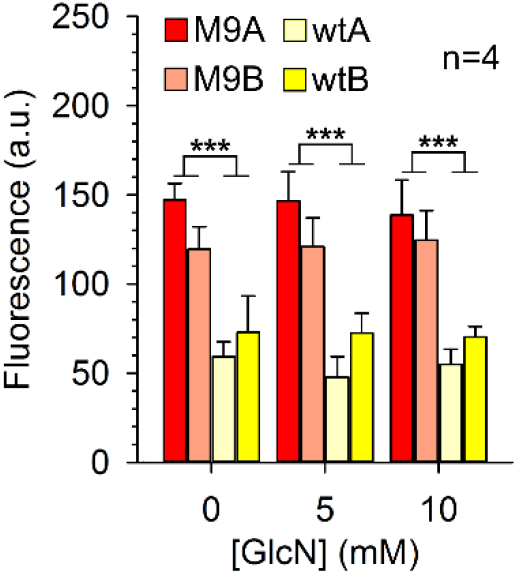
Constitutive *mCherry-glmS* ribozyme knock-down in *L. tarentolae* that was grown in BHI medium. The normalized mCherry fluorescence was determined for *L. tarentolae* with wild-type (wt) or mutant (M9) *mCherry-glmS* ribozyme. Two clonal cell lines (A and B) were analyzed for each construct. Parasites were grown in parallel in standard BHI medium in the presence of 0, 5 or 10 mM glucosamine (GlcN). All data points are the mean ± standard deviation from four biological replicates. Statistical analyses were performed in SigmaPlot 13 using the one-way ANOVA method (*** p < 0.001). Differences between cell lines A and B or between cultures with 0, 5 and 10 mM glucosamine were not significant (p > 0.05).

### Chromosomal *glmS*-tagging of *PF16, APRT* and *ERV*

In the final set of experiments, we used the CRISPR-Cas9 system for gene tagging in order to study a *glmS* ribozyme-dependent knock-down of chromosomal genes in *L. tarentolae*. As a proof-of-principle, the methodology was first tested for mCherry-tagged Pf16 and Erv using plasmid pPLOT-mCherry-puro^12^. C-terminally mCherry-tagged Pf16 was shown to localize to the flagellum (Fig. S2), thus confirming the suitability of the CRISPR-Cas9 method for gene tagging in *L. tarentolae*. In contrast, neither N-nor C-terminal mCherry-tagging of Erv resulted in viable parasites, suggesting that the tag interferes with an essential protein function. Next, we fused *PF16* (Fig. 6), *APRT* (Fig. 7) and *ERV* (Fig. 8) with the wild-type *glmS* ribozyme as well as the *glmS* M9 control. We therefore generated a template plasmid for PCR-based 3’-tagging of a gene of interest that is compatible with the LeishGEdit system^12,32^. Parasites with plasmid pTB007 were co-transfected with PCR products encoding (i) a sgRNA for the introduction of a double-strand break close to the stop codon of the gene of interest and (ii) an insert for a triple HA-tag, the *glmS* ribozyme and a resistance cassette for the repair of the double-strand break (Fig. S3). Following transfection and selection on BHI agar plates, colonies appeared after 7-14 days.

**Figure 6.**
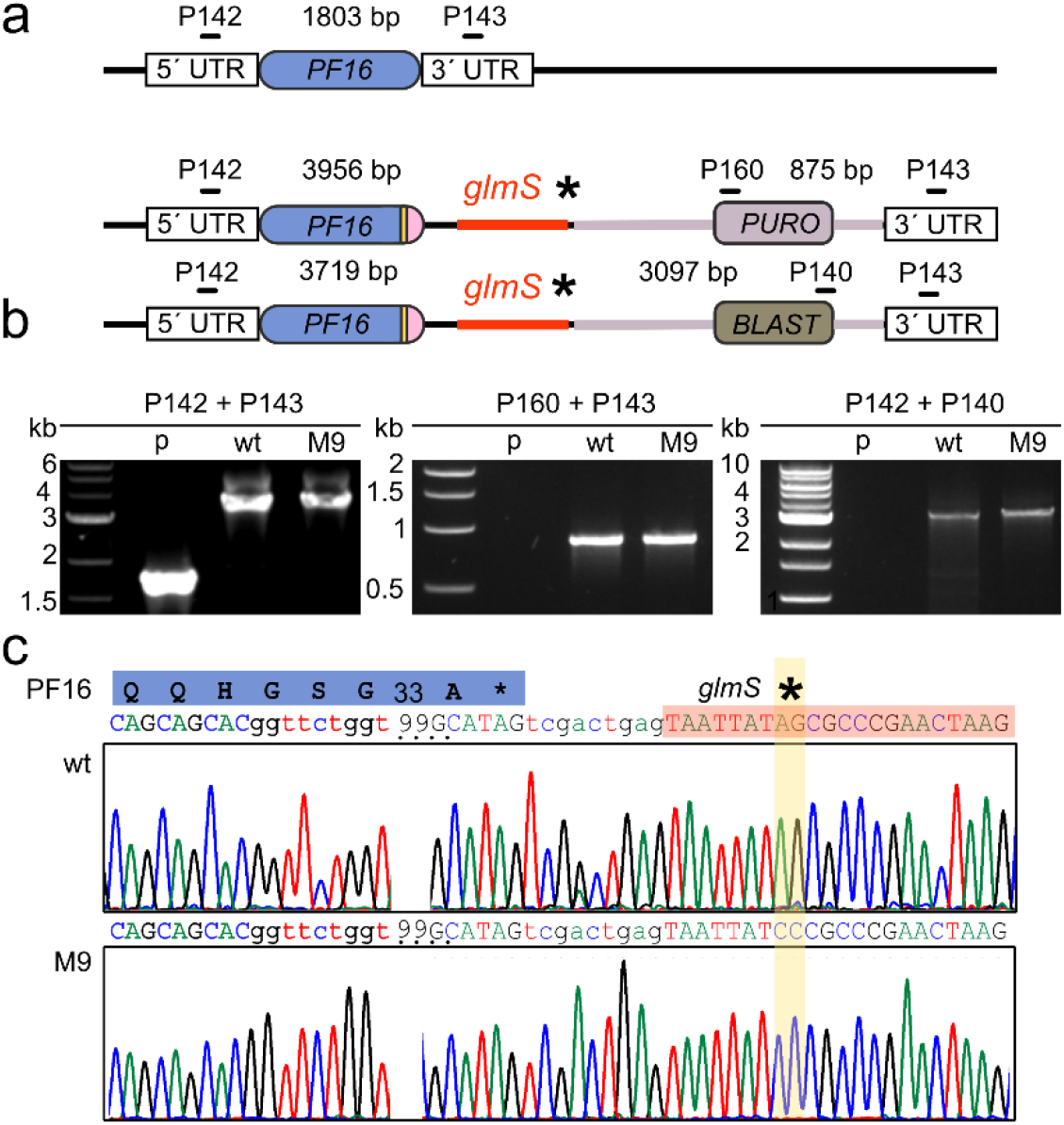
Generation and validation of chromosomal *glmS*-tagged *PF16* in *L. tarentolae* using the CRISPR-Cas9 system. (**a**) Schematic representation of 3’-tagged *PF16* after knock-in experiments with a cassette that encodes a triple HA-tag, the *glmS* ribozyme and puromycin *N*-acetyltransferase (*PURO*) or blasticidin deaminase (*BLAST*). Primer positions and expected product sizes from analytical PCR reactions are highlighted. (**b**) PCR analyses of genomic DNA of the parental strain (p) and of a wild-type (wt) or two M9 *glmS*-tagged cell lines after transfection and selection. PCR products were separated by agarose gel electrophoresis confirming the expected product sizes from panel a. Primer pairs are indicated on top of each gel. (**c**) Sequencing of PCR products from genomic DNA confirmed the tagging and the correct sequences for the wt or M9 *glmS* ribozyme.

**Figure 7.**
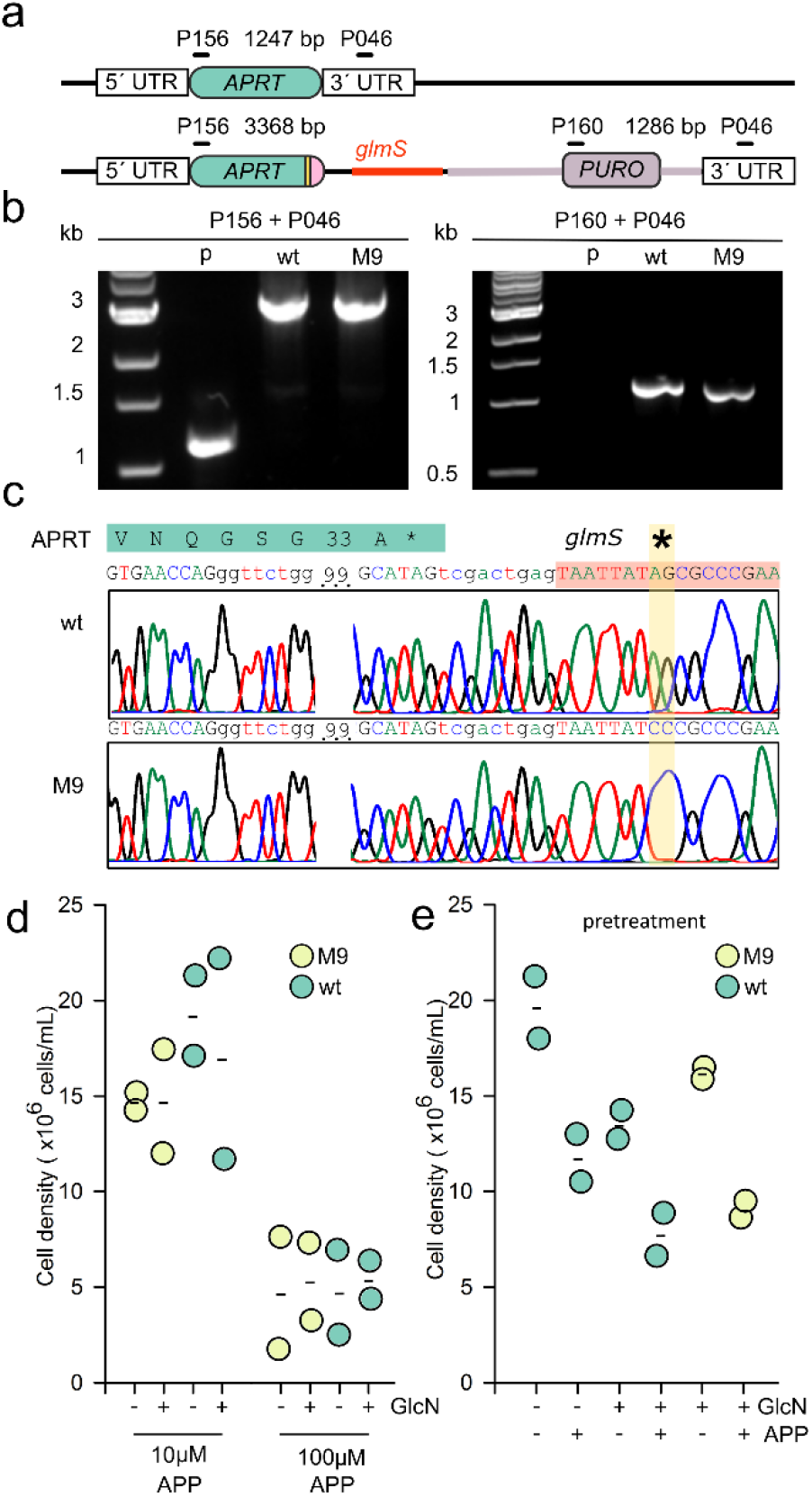
Generation and validation of chromosomal *glmS*-tagged *APRT* in *L. tarentolae* using the CRISPR-Cas9 system. (**a**) Schematic representation of 3’-tagged *APRT* after knock-in experiments with a cassette that encodes a triple HA-tag, the *glmS* ribozyme and puromycin *N*-acetyltransferase (*PURO*). Only one of both *L. tarentolae* gene copies is shown. Primer positions and expected product sizes from analytical PCR reactions are highlighted. (**b**) PCR analyses of genomic DNA of the parental strain (p) and of a wild-type (wt) or two M9 *glmS*-tagged cell lines after transfection and selection. PCR products were separated by agarose gel electrophoresis confirming the expected product sizes from panel a. Primer pairs are indicated on top of each gel. (**c**) Sequencing of PCR products from genomic DNA confirmed the tagging and the correct *APRT* fusion sequences for the wild-type (wt) or M9 *glmS* ribozyme. (**d**) Growth analysis of *APRT* lines with fused wt and inactive M9 riboswitches in supplemented MEM. Parasites were exposed for 72 h to 10 or 100 μM 4-aminopyrazolopyrimidine (APP) with or without 10 mM glucosamine (GlcN). (**e**) Cells were grown for two passages in supplemented MEM with or without 5 mM glucosamine as a pretreatment before the cultures were grown for 72 h with 10 μM APP and 5 mM glucosamine. Data from two independent biological replicates are shown.

**Figure 8.**
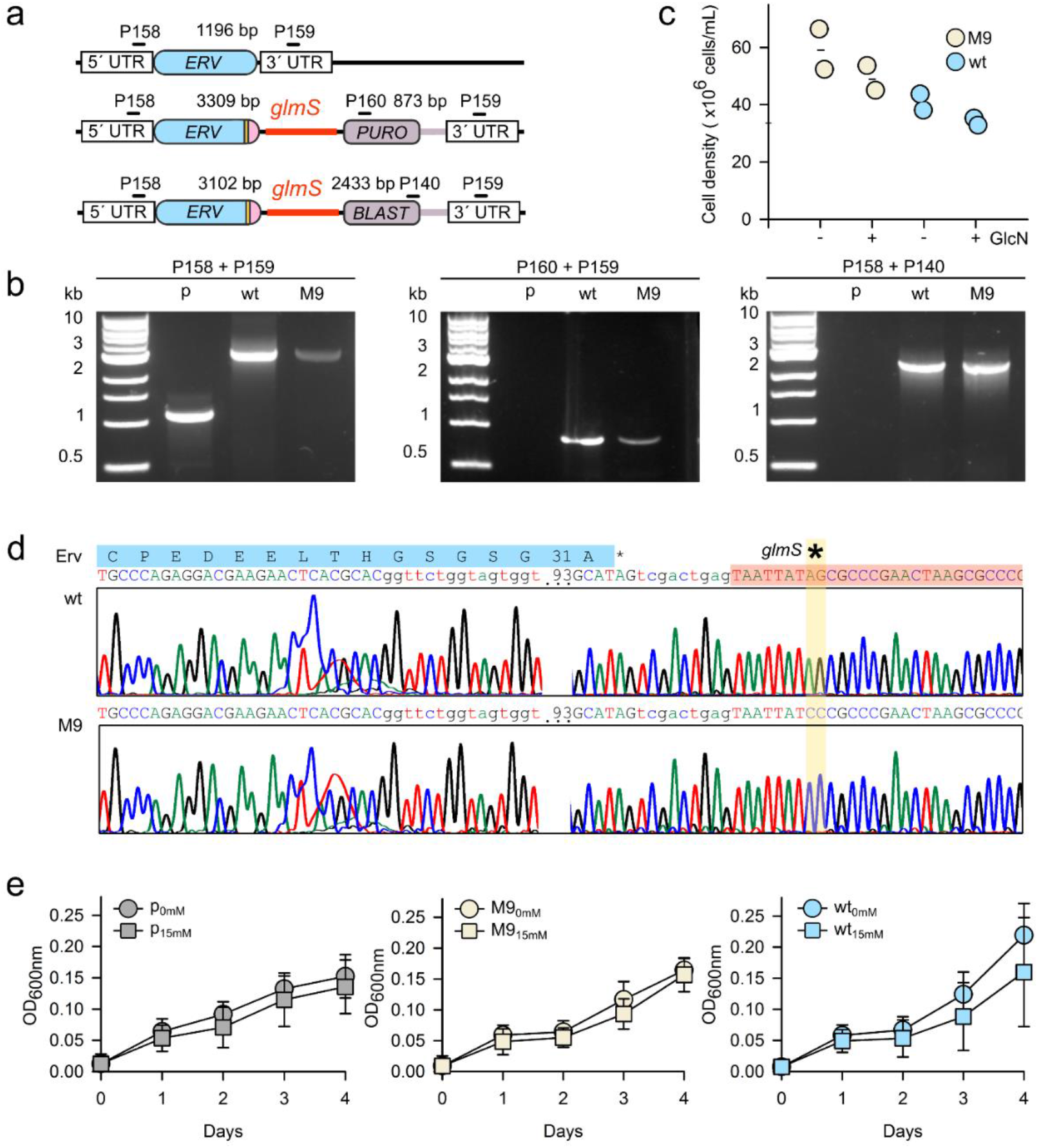
Generation and validation of chromosomal *glmS*-tagged *ERV* in *L. tarentolae* using the CRISPR-Cas9 system. (**a**) Schematic representation of 3’-tagged *ERV* after knock-in experiments with a cassette that encodes a triple HA-tag, the *glmS* ribozyme and puromycin *N*-acetyltransferase (*PURO*) or blasticidin deaminase (*BLAST*). Primer positions and expected product sizes from analytical PCR reactions are highlighted. (**b**) PCR analyses of genomic DNA of the parental strain with plasmid Tb007 (p) and of a wild-type (wt) or two M9 *glmS*-tagged cell lines after transfection and selection. PCR products were separated by agarose gel electrophoresis confirming the expected product sizes from panel a. Primer pairs are indicated on top of each gel. (**c**) Cell densities of strains with wt or M9 *glmS*-tagged *ERV* after 4 days continuous incubation in supplemented MEM with or without 15 mM glucosamine (GlcN). Data from two independent biological replicates are shown. (**d**) Sequencing of PCR products from genomic DNA confirmed the tagging and the correct sequences for the wt or M9 *glmS* ribozyme. (**e**) Growth curve analysis of the parental strain (p) as well as strains with wt or M9 *glmS*-tagged *ERV* in supplemented MEM with or without 15 mM GlcN. Values represent the mean ± standard deviation from three independent biological replicates. Statistical analysis using the one-way ANOVA method in SigmaPlot 13 did not reveal a significant difference among strains (p > 0.05).

Tagging of both *PF16* alleles required simultaneous selection with 20 μg/mL puromycin and 10 μg/mL blasticidin (Fig. 6a). PCR analyses confirmed the knock-in and replacement of untagged *PF16* (Fig. 6b). Sequencing of PCR products from genomic DNA revealed the correct sequences for wild-type *glmS* ribozyme and the *glmS* M9 control (Fig. 6c). However, all strains remained fully motile in the presence or absence of 10 mM glucosamine in supplemented MEM. Hence, the *glmS* knock-down system did not result in a comparable phenotype to the knock-out control in Fig. 2c.

Tagging of both *APRT* alleles with *glmS* was achieved after a single round of selection with 20 μg/mL puromycin (Fig. 7a). PCR analyses confirmed the expected modification of the *APRT* locus (Fig. 7b) and sequencing of PCR products from genomic DNA revealed the correct sequences for wild-type *glmS* ribozyme and the *glmS* M9 control (Fig. 7c). All strains remained sensitive towards 4-aminopyrazolopyrimidine in the presence or absence of 10 mM glucosamine in supplemented MEM, regardless whether wild-type *glmS* or the M9 control was fused to *APRT* (Fig. 7d). The susceptibilities of the *APRT* fusion constructs with wild-type *glmS* or the M9 control remained also highly similar when the cells were pretreated with or without glucosamine before the addition of 4-aminopyrazolopyrimidine (Fig. 7e). Thus, the knock-out phenotype for *APRT* in Fig. 2f could not be mimicked using the *glmS* knock-down system.

Tagging of both *ERV* alleles was achieved after simultaneous selection with 20 μg/mL puromycin and 10 μg/mL blasticidin (Fig. 8a). PCR analyses confirmed the knock-in and replacement of untagged *ERV* (Fig. 8b). However, tagging of *ERV* with the wild-type *glmS* ribozyme only slightly decreased cell growth in supplemented MEM as compared to the M9 control. The addition of 15 mM glucosamine appeared to have only a minor effect on both cell lines (Fig. 8c). Sequencing of PCR products from genomic DNA confirmed the expected sequences for wild-type *glmS* ribozyme and the *glmS* M9 control (Fig. 8d) and a more detailed growth analysis also revealed no significant growth defects for wild-type *glmS*-tagged *ERV* in the presence of up to 15 mM glucosamine (Fig. 8e). In summary, we showed that the LeishGEdit CRISPR-Cas9 system can be also applied to gene tagging and knock-in studies in *L. tarentolae* whereas the *glmS* ribozyme system appears to lack efficiency for general chromosomal knock-down studies in this organism.

## Discussion

The CRISPR-Cas9 system has been recently established for a variety of Leishmania parasites^10–13,32,40^. Here we demonstrated that a robust and simple PCR- and T7 RNA polymerase-based CRISPR-Cas9 system can be also used for the rapid and reliable generation of knock-out and knock-in cell lines in *L. tarentolae*. We successfully deleted the genes encoding the flagellar motility factor Pf16 and the salvage-pathway enzyme adenine phosphoribosyltransferase. The results confirmed their expected relevance for parasite motility and drug susceptibility, respectively^12,13,33^. Furthermore, using the CRISPR-Cas9 system we were unable to delete the *L. tarentolae* gene that encodes the mitochondrial flavoenzyme Erv in accordance with previous results that were obtained for *L. infantum* using traditional genetics^35^. Hence, two unrelated genetic studies in *Leishmania*, as well as RNAi experiments in *T. brucei*^38,41^, show that Erv is crucial for parasite survival. The results are encouraging for potential intervention strategies taking into account that parasite Erv homologues and human Erv/ALR have very different structures and also employ a deviating enzyme mechanism^37,41^.

Whilst the CRISPR-Cas9 system will facilitate the identification of essential genes in important human pathogens, the characterization of these genes and the suitability of *Leishmania* parasites as model organisms in the post-genomic era will probably depend on the development of alternative knock-down methods. To date, functional genomics in kinetoplastid parasites and the down-regulation of a gene of interest are restricted to only a few species because of a limited method repertoire^42^. For example, CRISPR interference^43^ would cause a simultaneous down-regulation of numerous genes in kinetoplastid parasites, which appear to lack gene-specific promoters and have head-to-tail oriented genes that are co-transcribed as polycistronic units^44,45^. Furthermore, while RNAi is commonly used in *Trypanosoma brucei*^18,22^ and was also shown to be functional in *L. braziliensis*^21,46^, attempts to implement RNAi in *T. cruzi, L. major* or *L. donovani* failed^47,48^. These and many other kinetoplastid parasites including *L. tarentolae* have lost key components of the RNAi machinery^21,22^. With the exception of the functional RNAi system in the *Leishmania* subgenus *Viannia*^21^, knock-down experiments in *Leishmania* rely so far on protein-destabilization strategies^49–52^ or tetracycline repressor systems^53,54^. Applications of these knock-down methods in human pathogens, as reported for *L. major, L. mexicana, L. braziliensis* and *L. donovani*, might have drawbacks, for example, because of protein-tagging or antibiotic toxicity. Interference with protein function could, for example, explain why we were unable to add an mCherry tag to *L. tarentolae* Erv in contrast to Pf16.

The *glmS* ribozyme system has become a valuable alternative knock-down method in eukaryotes since its discovery in 2004 (ref. ^26^). For example, the system is now commonly used in the apicomplexan malaria parasite *Plasmodium falciparum* for the analysis of drug targets as well as essential genes and processes^30,55–58^. A *glmS* ribozyme system has also been established for yeast and, very recently, for *T. brucei* and *T. cruzi*^31,59,60^. The *glmS* ribozyme system now offers the opportunity for medium-dependent constitutive or inducible knock-down experiments of episomal constructs in *L. tarentolae*. Although the knock-down efficiency for our episomal *mCherry-glmS* reporter was below 80%, the *glmS* M9 controls revealed that the inducible down-regulation with 10 mM glucosamine in supplemented MEM was slightly stronger than the constitutive down-regulation in standard BHI medium. Since GFP and mCherry are rather stable proteins, the down-regulation of other proteins might be more efficient. Higher glucosamine concentrations could be used for short-term knock-downs as long as potential glucosamine-dependent off-target effects are addressed with *glmS* M9 controls. Glucosamine toxicity is not a concern for the constitutive down-regulation in standard BHI medium, which could be explained by relatively high endogenous glucosamine-6-phosphate concentrations as suggested recently for the *glmS* ribozyme system in *T. cruzi*^60^. Under these conditions, episomal *glmS* M9 constructs only serve as controls in order to quantify the degree of down-regulation. Whether a *glmS* ribozyme-dependent down-regulation of episomal constructs is sufficient for phenotypic and functional analyses in *L. tarentolae* might depend on the protein of interest. Nevertheless, the simplicity of the constitutive knock-down system in standard BHI medium might facilitate the generation and phenotypic screening of episomal knock-down libraries. Regarding chromosomal *glmS* knock-down studies, tagging of three different genes did not result in the expected phenotypes (as shown for our characterized knock-out strains). Thus, the *glmS* system has intrinsic limitations that impede its general application for chromosomal knock-down studies in *L. tarentolae*. In our opinion, the *glmS* ribozyme system might give similar results in other *Leishmania* species despite the presence of *N*-acetyl glucosamine 6-phosphate deacetylase, which was recently suggested to be one key factor for the functionality of the system in kinetoplastid parasites^60^. Whether the suitability of the *glmS* ribozyme system in *Leishmania* depends on (invariable) biological factors in addition to the medium composition remains to be addressed in future studies.

In conclusion, the CRISPR-Cas9 system facilitates the genetic manipulation of *L. tarentolae* and provides a powerful technology for future analyses of this nonpathogenic model organism. In contrast, the *glmS* system failed to produce a detectable phenotype for chromosomal knock-downs but might be used for medium-dependent constitutive or inducible episomal knock-downs in *L. tarentolae*. The development of an unbiased and highly efficient knock-down system for important pathogens and model systems remains one of the biggest challenges in *Leishmania* molecular biology.

## Materials and Methods

### Generation of *L. tarentolae* CRISPR-Cas9 knock-out cell lines

*L. tarentolae* promastigote mutants were obtained by reproducing the CRISPR-Cas9 system reported by Beneke et al. using plasmids pTPuro, pTBlast and pTB007^12,13^. Donor DNA for the repair of double-strand breaks (containing a selectable marker and 30 nt homology arms) as well sgRNA templates for the excision of *PF16, APRT* or *ERV* were generated according to the LeishGEdit method^12,32^. Primer sequences for the generation of sgRNA templates and the amplification of targeting cassettes for strain *L. tarentolae* ParrotTarII were obtained online (www.leishgedit.net) and are listed in Supplementary Tables 1–4. Puromycin or blasticidin resistance cassettes with 5’- and 3’-homology regions for the repair of excised *PF16, APRT* or *ERV* were amplified by PCR using plasmid pTPuro or pTBlast. Transfections were performed as outlined below in a parental cell line that transiently expressed Cas9 nuclease and T7 RNA polymerase from plasmid pTB007.

### Generation of *GFP*- and *mCherry-glmS* reporter constructs

Plasmids encoding GFP- or mCherry-reporters were cloned using the primers that are listed in Supplementary Table 5. To clone reporter construct pX-*GFP-glmS*, the fusion construct *GFP-glmS* was PCR-amplified using primers 114 and 115 with vector pARL-*GFP-glmS* as a template. The PCR product was subsequently cloned into the *Bam*HI and *HindIII* restriction sites of vector pX (ref.^61^). To generate pX-*mCherry-glmS, mCherry* was first PCR-amplified from vector pLEXY_IE-blecherry4 (JenaBioscience) using primers 127 and 128 and cloned into pCR 2.1 TOPO (Thermo Fisher Scientific) to generate pCR 2.1 TOPO-*mCherry*. The M9 mutation reported by Winkler et al.^26^ (*glmS*^M9^) was introduced by PCR using primer 129 with the desired mismatch, antisense primer 130 and vector pARL-*GFP-glmS* as a template. The wild-type *glmS* ribozyme-encoding sequence (*glmS*^wt^) was PCR-amplified from vector pARL-*GFP-glmS* using primers 131 and 130. The *glmS*^wt^ and *glmS*^M9^ PCR products were cloned into the *Bsr*GI and *Hin*dIII restriction sites of pCR 2.1 TOPO-*mCherry* to generate pCR 2.1 TOPO-*mCherry-glmS*^wt^ and pCR 2.1 TOPO-*mCherry-glmS*^M9^, respectively. The inserts *mCherry-glmS*^wt^ and *mCherry-glmS*^M9^ were subsequently excised from the according pCR 2.1 TOPO constructs and subcloned into the *Bam*HI and *HindIII* restriction sites of pX. All constructs were confirmed by Sanger sequencing (SEQ-IT GmbH & Co. KG). The sequences for *GFP-glmS, mCherry-glmS*^wt^ and *mCherry-glmS*^M9^ are listed in the Supplementary material.

### Generation of chromosomally tagged *L. tarentolae* cell lines

Constructs for chromosomal *mCherry*-tagging of *PF16* or *ERV* were PCR-amplified using plasmid pPLOT-mCherry-puro as a template^12,32^. For chromosomal *glmS*-tagging of *PF16, APRT* or *ERV*, we generated plasmids that are compatible with the LeishGEdit system using the primers that are listed in Supplementary Table 6. We therefore modified plasmids pMOTag-glmS-4H wt and M9 from ref.^59^ (addgene plasmids #106378 and #106379, respectively) so that they contain the puromycin or blasticidin cassette with primer binding sites 4 and 5 from the LeishGEdit system. To generate these plasmids, (i) we amplified the 3xHA-*glmS* wt and M9 sequences from pMOTag-glmS-4H wt and M9 by PCR with primer 173, which contains the *ApaI* restriction site and the primer binding site 5, and primer 174, which contains an *NcoI* restriction site. (ii) Resistance marker cassettes against puromycin or blasticidin suitable for 3’-tagging were amplified by PCR from plasmids pTPuro or pTBlast with primers 175 and 176 containing *Nco*I and *Bam*HI restriction sites, respectively. (iii) Fragments generated from steps (i) and (ii) were ligated using the *Nco*I restriction site and finally subcloned into the pMOTag-glmS-4H backbone using the *Apa*I and *Bam*HI restriction sites. The resulting plasmids served as templates to generate the donor DNA for the repair of double-strand breaks in combination with the sgRNA templates for *PF16, APRT* or *ERV* as described above. The sequences of the the 3xHA-*glmS-Puro/Blast* cassettes are listed in the Supplementary material.

### Standard *L. tarentolae* culture and screen of alternative culture media

BHI powder (#237200) was purchased from BD Bioscience, AlbuMax II lipid-rich BSA, heat inactivated FBS as well as DMEM (#42430), DMEM/F-12 (#11330), RPMI-1640 (#52400), IMDM (#12440), MEM (#31095) and McCoy’s 5A medium (#26600) were from Thermo Fisher Scientific. A 2 mg/mL stock solution of hemin chloride from Calbiochem was prepared in 0.05 M NaOH, sterile filtered (0.2 μm) and stored at −20°C. *L. tarentolae* promastigotes were cultured in 25 cm^2^ culture flasks under standard conditions at 27°C in BHI medium containing 37 mg/mL BHI powder and 10 μg/mL hemin as described previously^62^. MEM and McCoy’s 5A medium were supplemented with 25 mM HEPES and sterile filtered (0.2 μm) prior to use. Except for BHI medium, all media were supplemented with either 0.5% (w/v) AlbuMax II or 0%, 5% or 10% (v/v) FBS. The media were also supplemented with folic acid (0 μg/mL, 10 μg/mL or 50 μg/mL) and hemin chloride solution (5 μg/mL, 10 μg/mL or 20 μg/mL) resulting in 216 different test conditions that were compared with standard BHI medium as a control. *L. tarentolae* parasites from standard cultures were centrifuged in the mid-logarithmic phase at 1500 × *g* for 10 min at room temperature and washed in the different media without any supplements. After centrifugation, cells were resuspended in the according supplemented medium and diluted to an initial concentration of 5 × 10^6^ cells/mL. Technical triplicates of the test cultures were transferred to 48-well plates (500 μL per well) and incubated at 27°C for three days. Parasite growth and morphology were qualitatively evaluated every 24 h by standard and differential interference contrast (DIC) light microscopy. Based on the screen in 48-well plates, growth curves were subsequently determined in 25 cm^2^ T-flasks for the following media containing 10 mg/mL hemin chloride: IMDM supplemented with 5% FBS, 10% FBS or 0.5% AlbuMax™, MEM supplemented with 25 mM HEPES and 5% or 10% FBS, McCoy’s 5A Medium supplemented with 25 mM HEPES and 5% or 10% FBS, and standard BHI medium as a control. *L. tarentolae* parasites from standard cultures were centrifuged in the mid-logarithmic phase at 1500 × *g* for 10 min at room temperature and washed once with one of the supplemented media. After centrifugation, cells were resuspended in the according supplemented medium and diluted in a total volume of 10 mL to an initial concentration of 5 × 10^6^ cells/mL. The parasites were cultured at 27°C on a Rotamax 120 shaker at 50 rpm. Aliquots (100 μL) were removed every 24 h and fixated with paraformaldehyde before the cell density was determined with a Neubauer chamber. In order to avoid a putative nutrient depletion, the cultures were subsequently centrifuged at 1500 × *g* for 10 min at room temperature and the cells were resuspended in 10 mL fresh medium. The parasite morphology was documented for cultures in the mid-logarithmic phase on a microscope slide that was sealed with paraffin using a Zeiss LSM780 microscope and the software ZEN2010.

### Transfection and selection of *L. tarentolae*

For transfection, 10^7^ *L. tarentolae* promastigotes in mid-logarithmic phase were centrifuged at 1500 × *g* for 3 min at room temperature and washed with 1 mL transfection buffer (21 mM HEPES, 137 mM NaCl, 5 mM KCl, 0.7 mM NaH_2_PO_4_, 6 mM glucose, pH 7.4). Parasites were subsequently resuspended in 100 μL nucleofector solution of the Basic Parasite Nucleofector Kit 2 (Lonza), combined with 50 μL plasmid solution containing 5-10 μg DNA, and pulsed in a Lonza Nucleofactor IIb using program U-033. Transfected parasites were allowed to recover in 10 mL standard BHI medium for 24 h before parasites were centrifuged and plated on BHI medium that was supplemented with 0.8% (w/v) agar, 0.08% (w/v) folic acid, 10% (v/v) FBS, 20 μg/mL hemin chloride and 40 μg/mL G418 disulfate (Calbiochem). Single colonies appeared 7-10 days after transfection and clonal parasites were subsequently transferred to liquid media. Targeting cassettes and sgRNA templates from 40 μL unpurified PCR product were transfected into 10^7^ parasites as described previously^12,63^ using the Lonza Nucleofactor IIb program X-001. Electroporated cells were allowed to recover in 2mL BHI medium without antibiotics for 16 h before being plated on BHI agar to obtain isogenic lines as described above. Single colonies appeared 7-14 days after transfection. The genome modifications were confirmed by analytical PCR using the primers in Supplementary Table 7 and by Sanger sequencing (SEQ-IT GmbH & Co. KG) of PCR products with genomic DNA as a template.

### Fluorescence measurements

Stock solutions of 500 mM D-(+)-glucosamine (#G4875 Sigma) were prepared in BHI medium or MEM. The pH was adjusted to 7.5 with 5 M NaOH before the stock solutions were sterile filtered (0.22 μm) and stored at −20 °C. In order to evaluate the *glmS* ribozyme-dependent down-regulation of the GFP reporter, clonal *L. tarentolae* cell lines with vector pX-*GFP-glmS* were grown at 27°C in standard BHI medium that was supplemented with 0.1 mg/mL G418 disulfate. Cultures (10 mL) in 25 cm^2^ T-flasks were diluted to an initial density of 5 × 10^6^ cells/mL, supplemented with 0 or 10 mM glucosamine and incubated for 24 h. Around 3 × 10^7^ cells were harvested by centrifugation at 1500 × *g* for 5 min and used for further fluorescence measurements as outlined below. To assess the *glmS* ribozyme-dependent down-regulation of the mCherry reporter, clonal *L. tarentolae* cell lines with vector pX-*mCherry-glmS*^wt^ or pX-*mCherry-glmS*^M9^ were grown at 27°C either in MEM that was supplemented with 10 μg/mL hemin, 5% (v/v) FBS, 25 mM HEPES, pH 7.4 and 0.1 mg/mL G418 disulfate or in standard BHI medium with 0.1 mg/mL G418 disulfate. Cultures (10 mL) in 25 cm^2^ T-flasks were diluted to an initial density of 5 × 10^6^ cells/mL or an OD_600nm_ of 0.2, supplemented with 0, 5 or 10 mM glucosamine and incubated for 24 h. Every 24 h, parasites were diluted to an OD_600nm_ of 0.2 in fresh supplemented MEM media with 0, 5 or 10 mM glucosamine. Three days after inoculation, cultures were harvested at 1500 × *g* for 5 min. The fluorescence from reporter cell lines was determined in a CLARIOstar fluorescence plate reader (BMG Labtech). Parasites were resuspended in 400 μL 100 mM MES/Tris buffer pH 6.0 to a final OD_600nm_ of 1.5. Technical duplicates of the suspensions (190 μL each) were transferred to a flat-bottom 96-well microplate (#353219, BD Biosciences). The microplate was centrifuged for 5 min at 30 × *g* and the fluorescence was subsequently measured with the following setting: GFP: λ_Exc_ = 485-15, dichroic 497.8, λ_Emis_ = 530-20, N° cycles 10, flashes/well 40, cycle time 6 s; mCherry: λ_Exc_ = 583-15, dichroic 602, λ_Emis_ = 623,5-20, N° cycles 6, flashes/well 40, cycle time 5 s. The data was averaged from three or four biological replicates as indicated. Statistical analyses were performed in SigmaPlot 13 using the one-way ANOVA method.

## Acknowledgements

G.L.T. was funded by the German Academic Exchange Service (DAAD). This work was in part funded by the DFG (grants DE 1431/9-1, DE 1431/10-1 and DE 1431/10-2 to M.D.). We thank Eva Gluenz for plasmids pTPuro, pTBlast, pPLOT-mCherry-puro and pTB007, Roberto Docampo for plasmids pMOTag-*glmS-4H* wt and M9, Michael Lanzer for plasmid pARL-*GFP-glmS*, Simone Eggert, Jessica Kehrer, Katharina Quadt, Carolina Andrade, Stefan Kins and Freddy Frischknecht for help with microscopy, and Prince Saforo Amponsah and Bruce Morgan for help with the CLARIOstar measurements.

## Author contributions

G.L.T., L.L. and M.D. designed the project and experiments. G.L.T. performed the knock-out and knock-down studies as well as CLARIOstar measurements. L.S. performed the *ERV* knock-down studies. L.L. performed the growth analyses in alternative media. G.L.T., L.S. and L.L. analyzed the data. M.D. supervised the study and wrote the manuscript. All authors edited and approved the manuscript.

## Supplementary Material and Methods

**Supplementary Figure 1.**
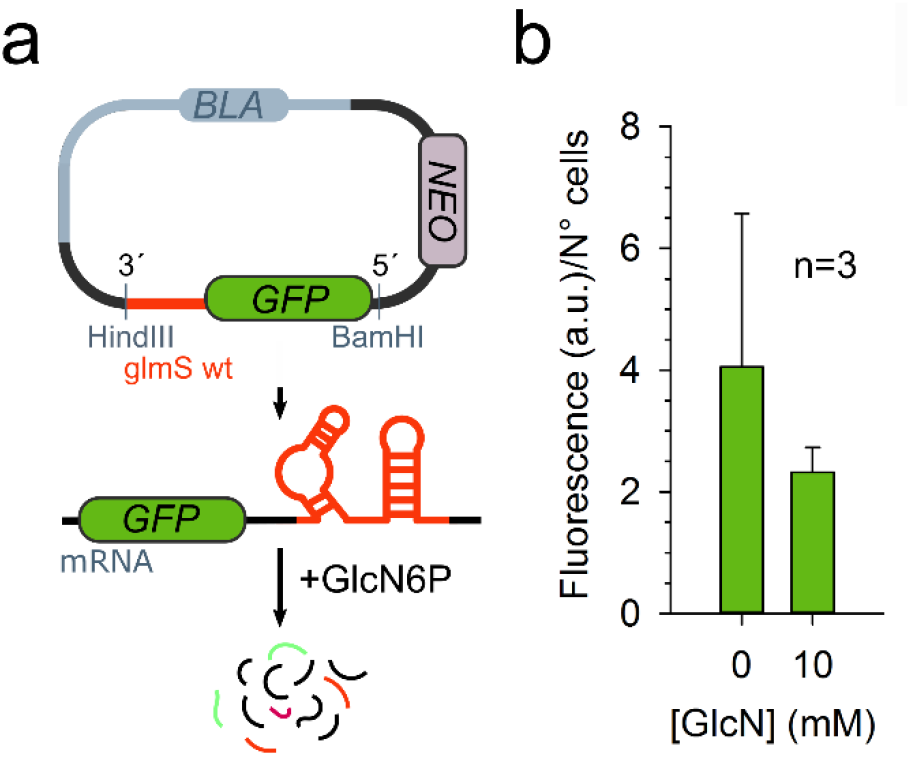
Initial *GFP-glmS* ribozyme knock-down attempts in BHI medium. (**a**) Schematic overview of the used *GFP-glmS* ribozyme construct in vector pX. The encoded mRNA should be cleaved and subsequently degraded in the presence of the activating ligand glucosamine-6-phosphate (GlcN6P). Selection markers against β-lactam antibiotics (*BLA*) and the aminoglycoside G418 (*NEO*) are indicated. (**b**) Normalized fluorescence from three biological replicates of a clonal *L. tarentolae* cell line that carried the *GFP-glmS* plasmid and that was grown in standard BHI medium with or without 10 mM glucosamine (GlcN).

**Supplementary Figure 2.**
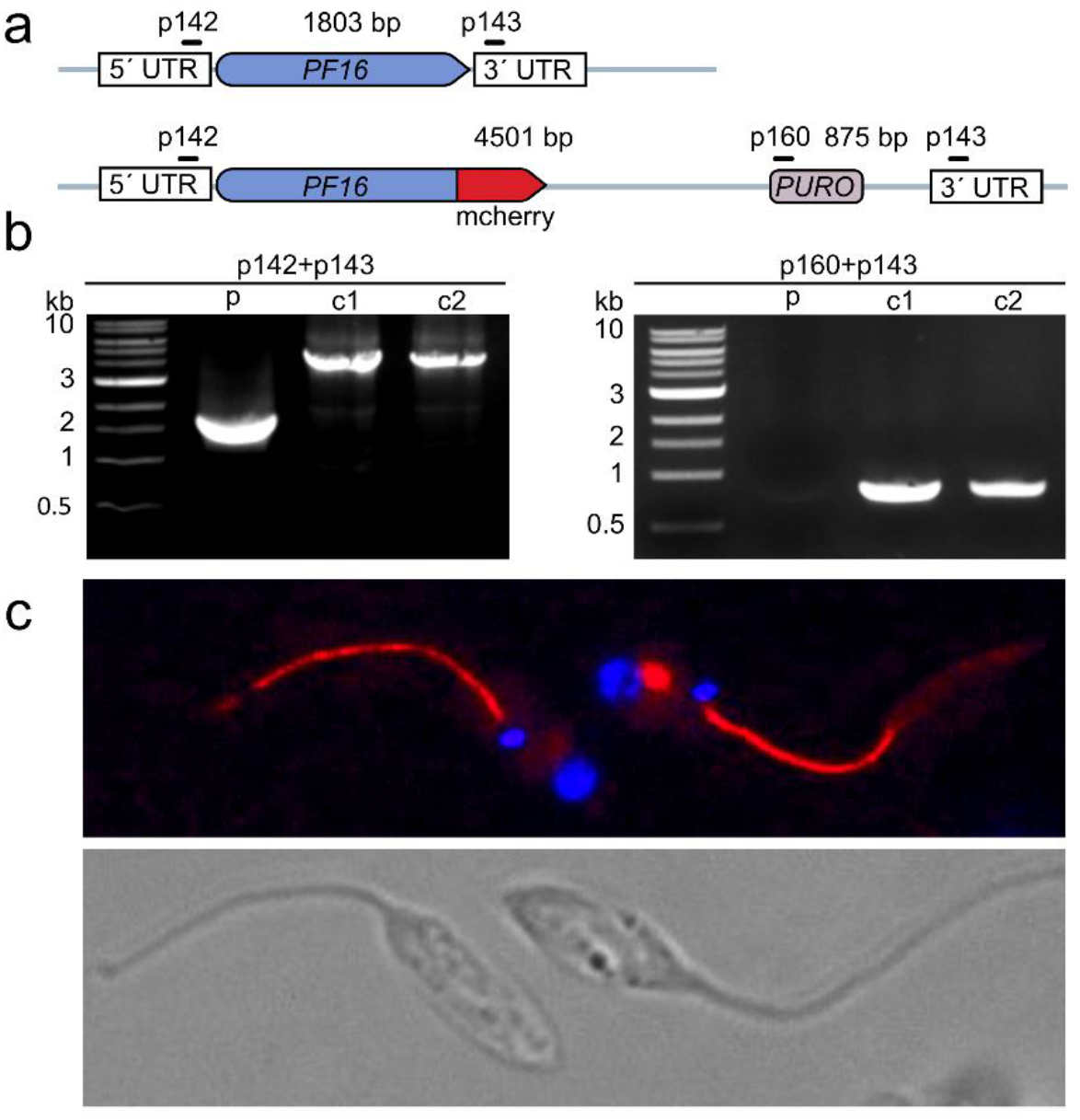
Generation and validation of chromosomal *mCherry*-tagging of *PF16* in *L. tarentolae* using the CRISPR-Cas9 system. (**a**) Schematic representation of 3’-tagged *PF16* after knock-in experiments with a cassette that encodes mCherry and puromycin *N*-acetyltransferase (*PURO*). Primer positions and expected product sizes from analytical PCR reactions are highlighted. (**b**) PCR analyses of genomic DNA of the parental strain (p) and of two clonal cell lines after transfection and selection. PCR products were separated by agarose gel electrophoresis and confirmed the expected product sizes from panel a. Primer pairs are indicated on top of each gel. (**c**) Fluorescence microscopy of *L. tarentolae* promastigotes with C-terminally mCherry-tagged Pf16.

**Supplementary Figure 3.**
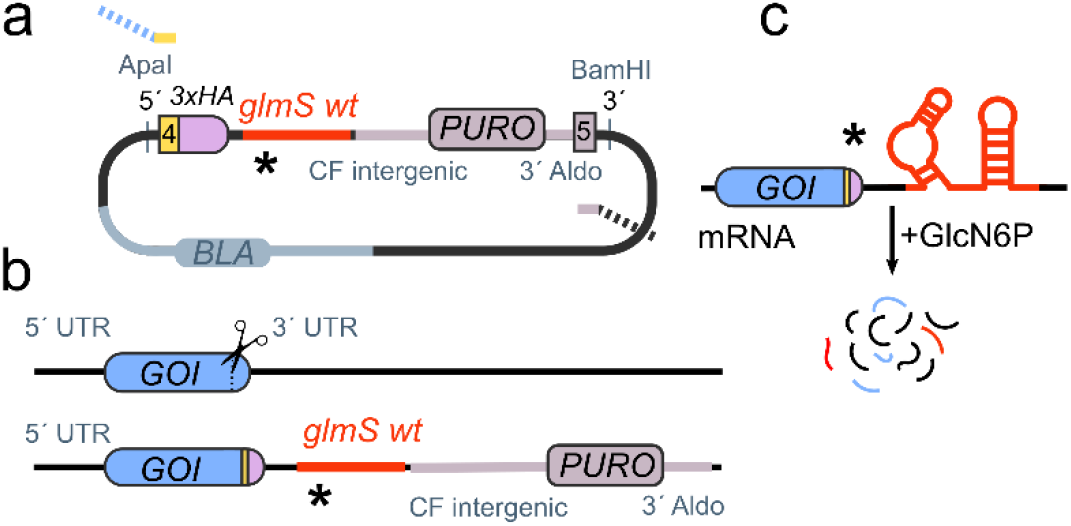
Generation of chromosomal *glmS*-tagged mutants. (**a**) Schematic representation of the *glmS* cassette within the template plasmid that is amplified by PCR (using primer sites 4 and 5 that are compatible with the LeishGEdit system^12,32^) to generate an insert for homologous recombination. The insert encodes a GSGSGSGSGS-linker that is fused to a triple HA-tag and a stop codon followed by the wild-type or M9 *glmS* sequence and a puromycin resistance cassette (*PURO*). (**b**) Strategy for 3’-tagging of a gene of interest (*GOI*) by inducing a sgRNA-targeted double-strand break that is repaired by homologous recombination using the insert from panel a. (**c**) Expected destabilization of the mRNA by the *glmS* ribozyme in the presence of the activating ligand glucosamine-6-phosphate (GlcN6P).

**Supplementary Table 1.**
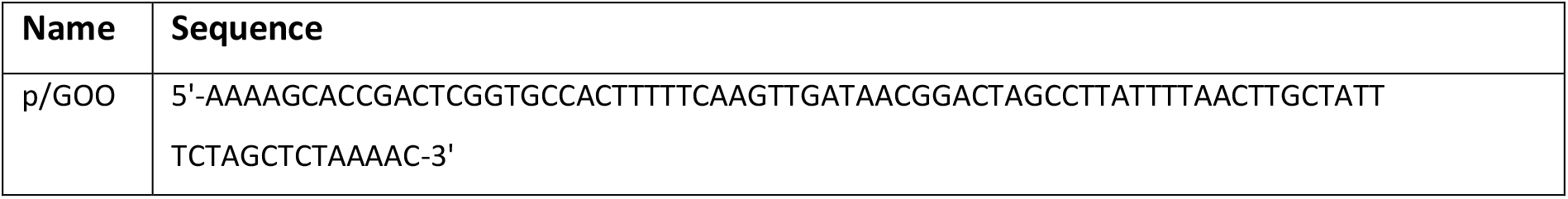
Universal sgRNA antisense primer.

**Supplementary Table 2.**
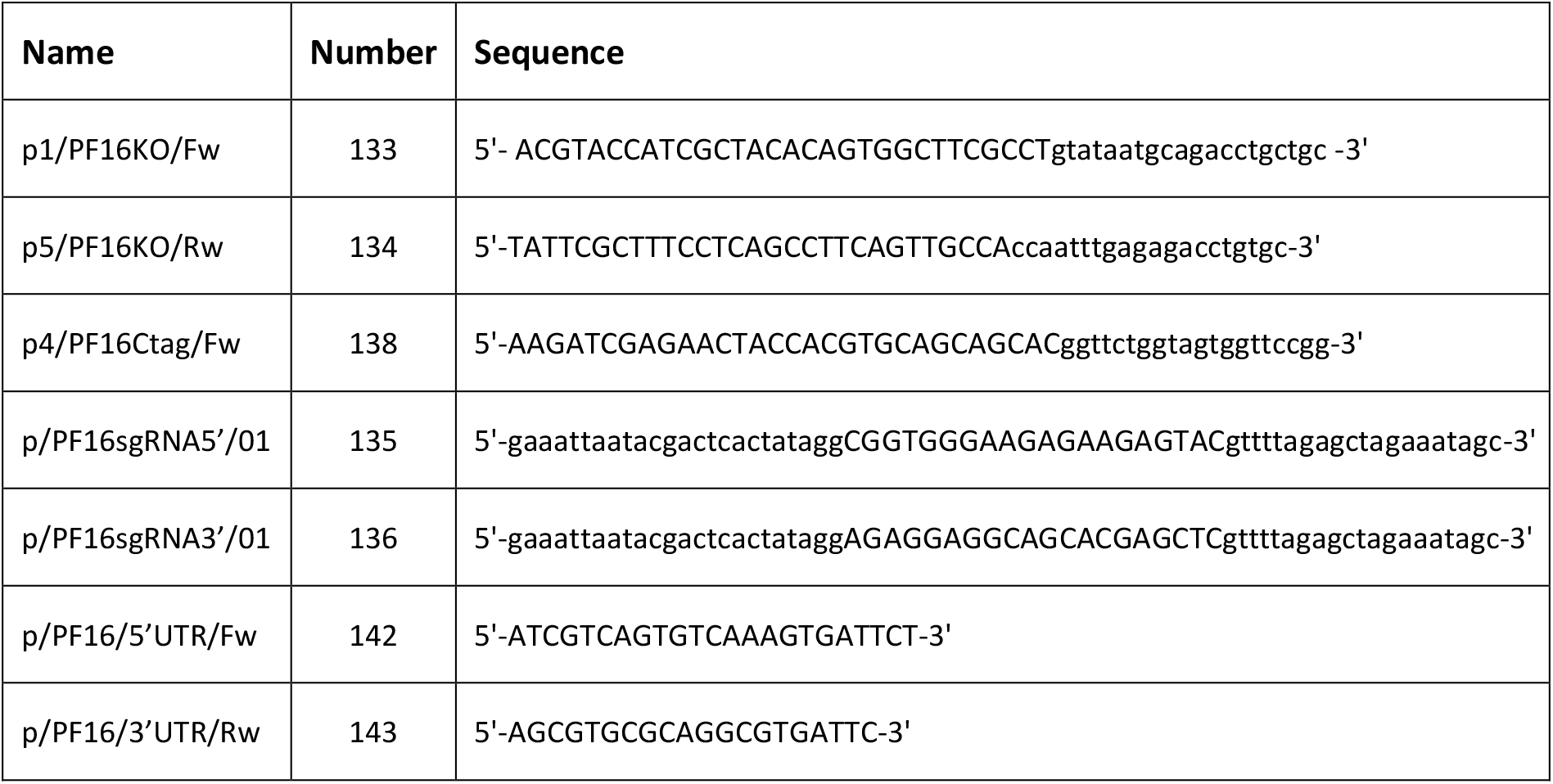
Knock-out and tagging of *PF16*. List of primers used to generate targeting cassettes, sgRNA guides and genotyping primers for knock out, mcherry and glmS tagging of *PF16*.

**Supplementary Table 3.**
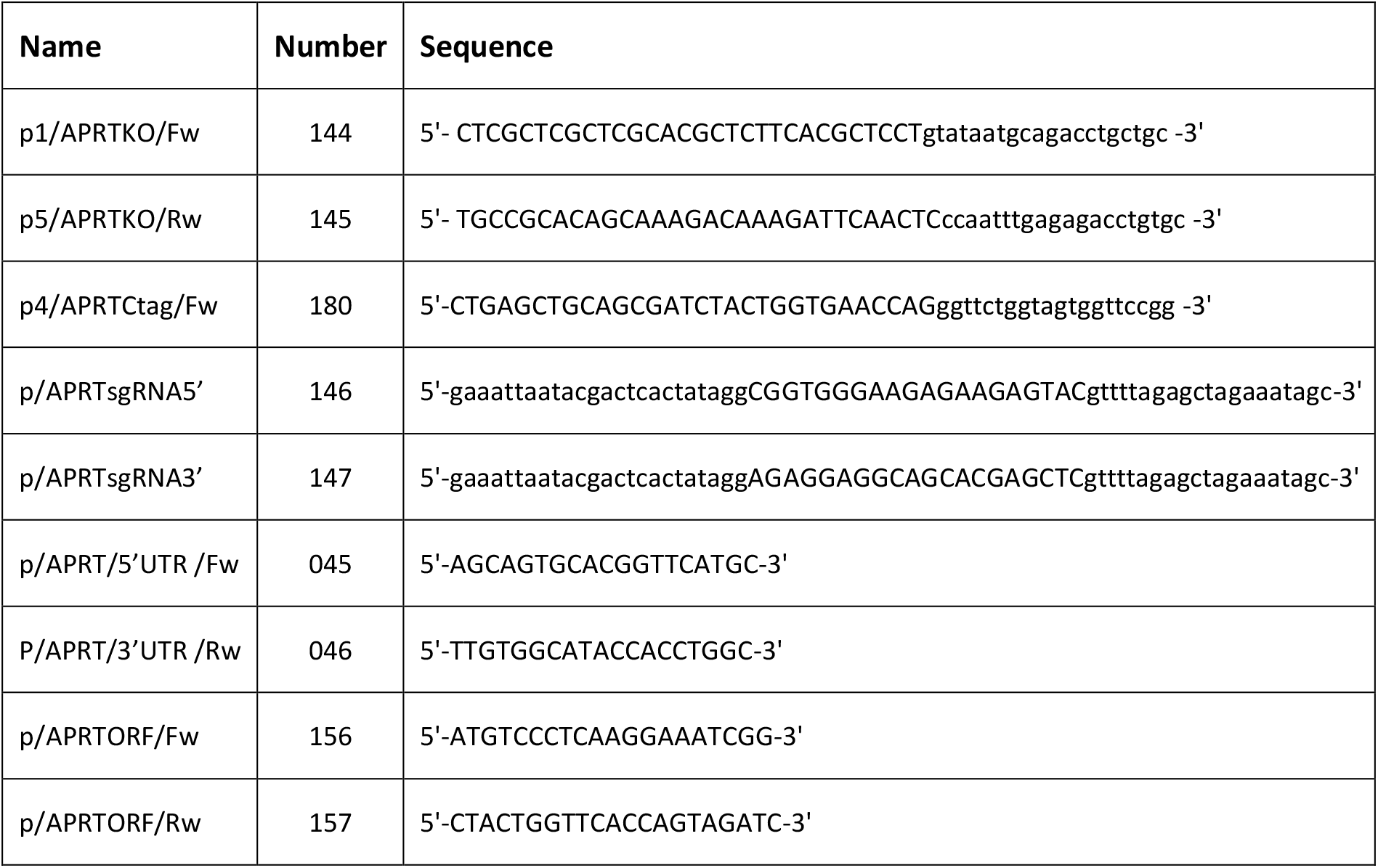
Knock-out and tagging of *APRT*. List of primers used to generate targeting cassettes, sgRNA guides and genotyping for knock out and glmS tagging of *APRT*.

**Supplementary Table 4.**
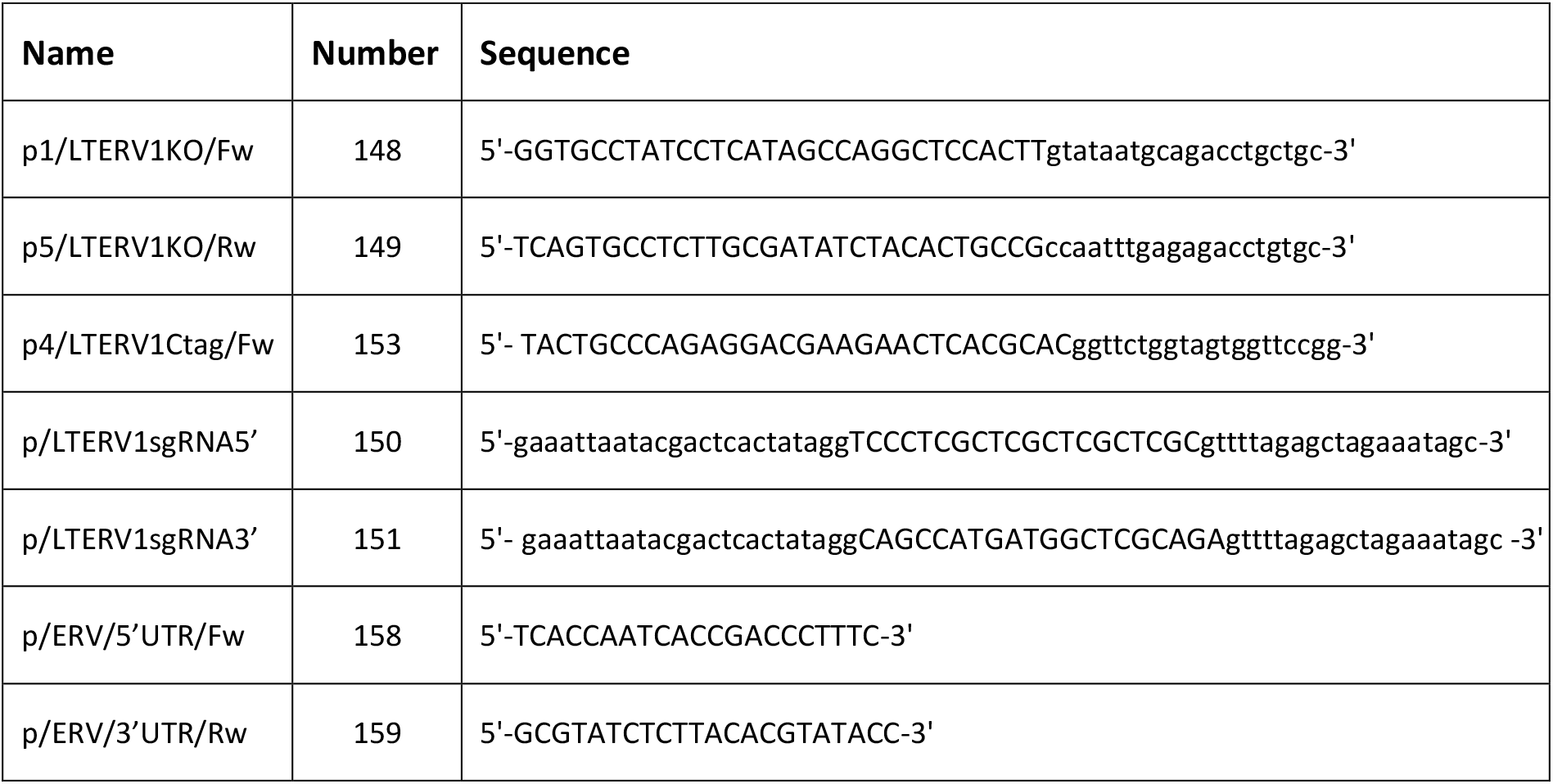
Knock-out and tagging of *ERV*. List of primers used to generate targeting cassettes, sgRNA guides and genotyping primer for knock out and glmS tagging of *ERV*.

**Supplementary Table 5.**
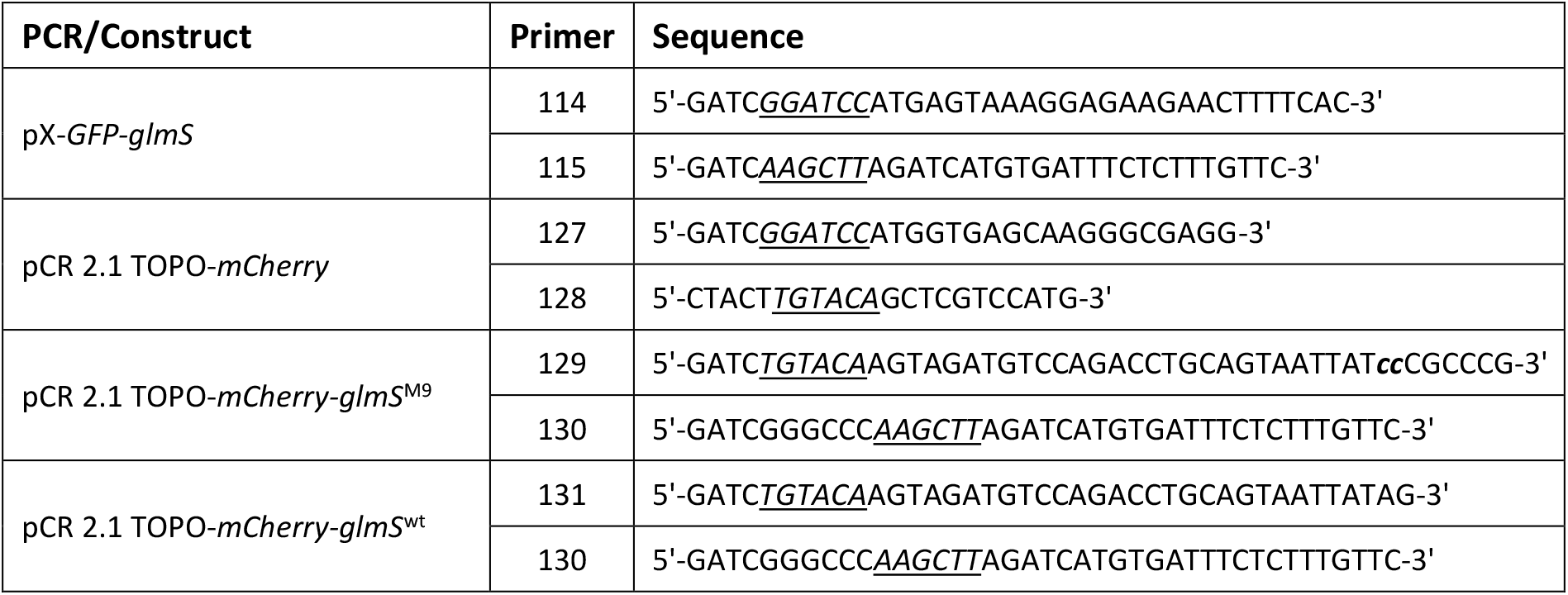
List of primers used for cloning of *glmS* reporter constructs. Restriction sites are underlined and mutated nucleotides are bold and italicized. See materials and methods for details.

**Supplementary Table 6.**
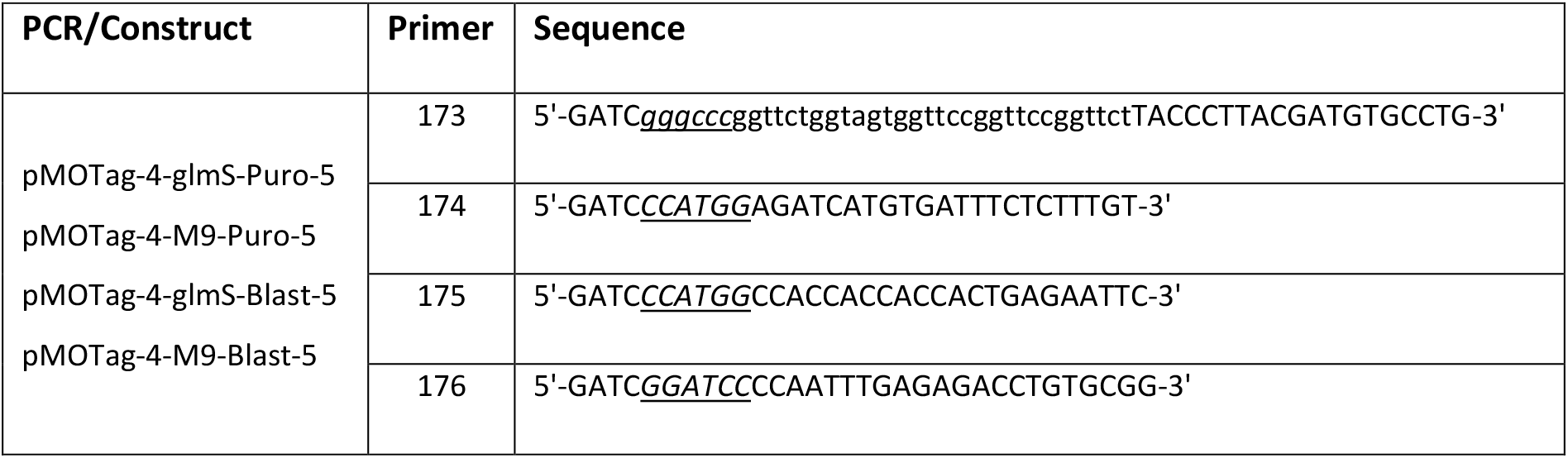
List of primers used for cloning of pMOTag constructs that are compatible with the LeishGEdit syste. Restriction sites are underlined and italicized. See materials and methods for details.

**Supplementary Table 7.**
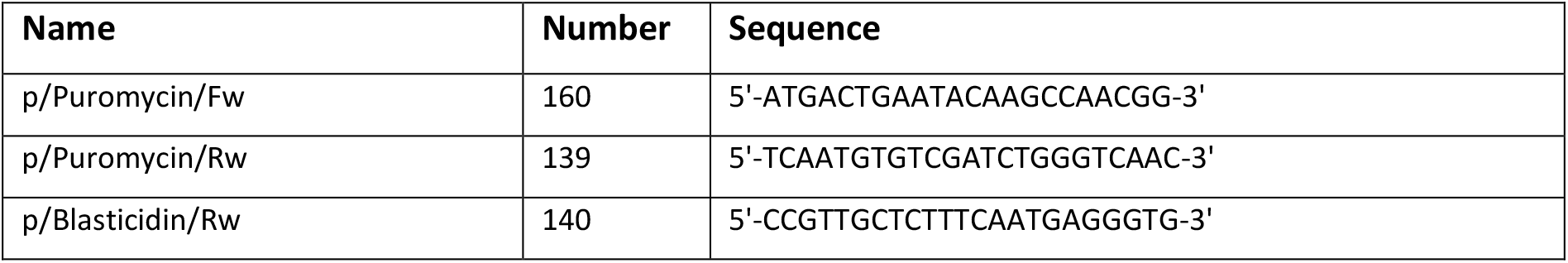
General genotyping primers.

## Sequence of *GFP-glmS* construct

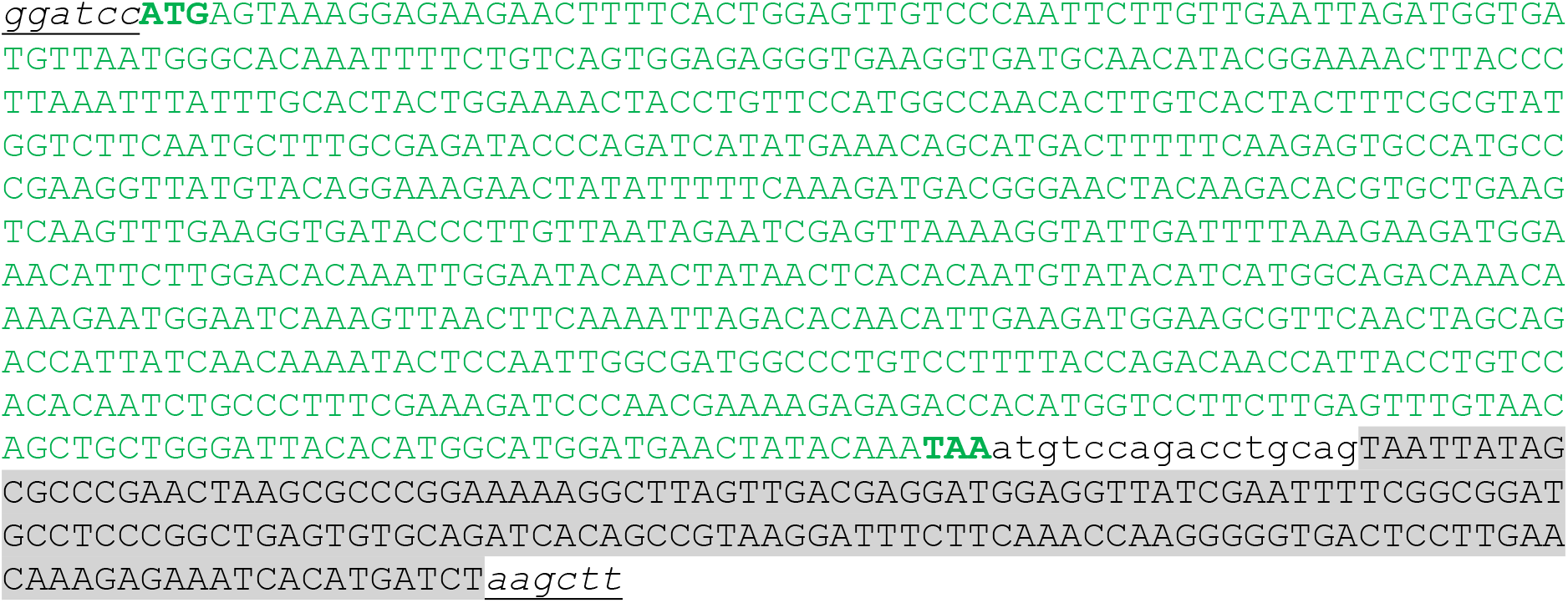

## Sequence of *mCherry-glmS*^wt^ construct

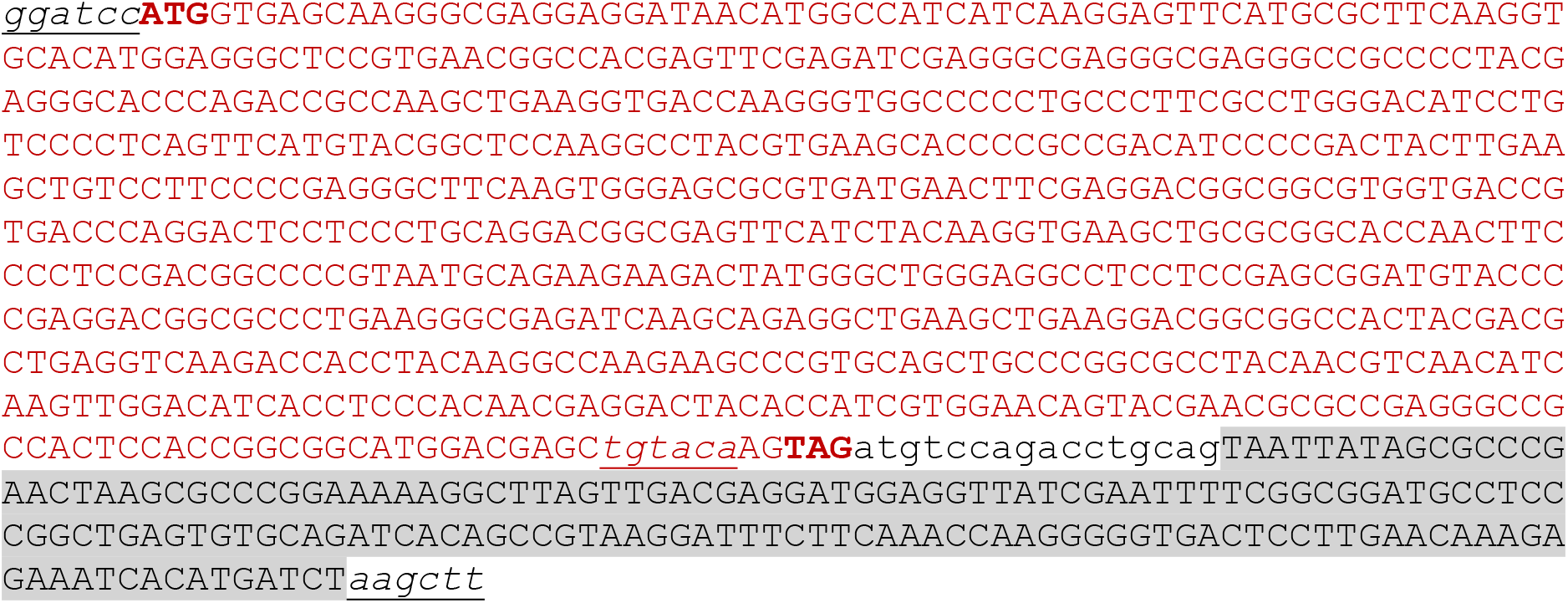

## Sequence of *mCherry-glmS*^M9^ construct

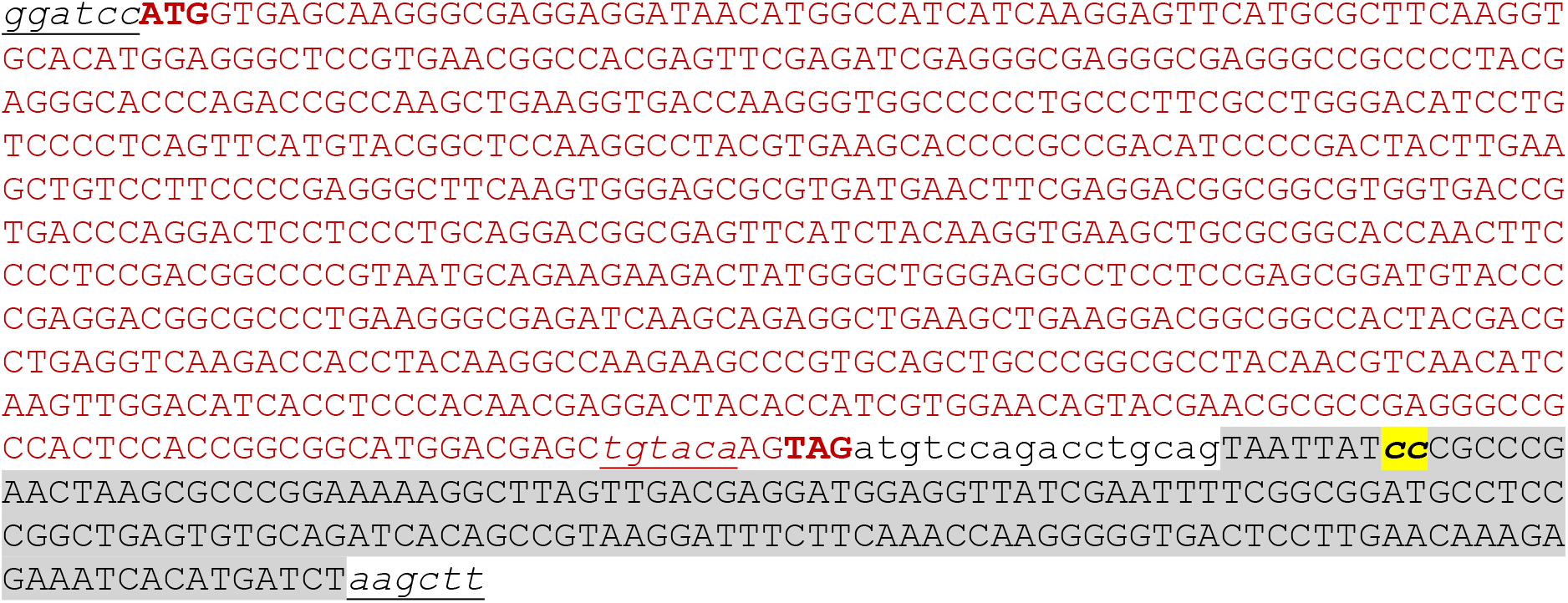

## Sequence of the *glmS*-tagging cassette in pMOTag-4-glmS-puro-5

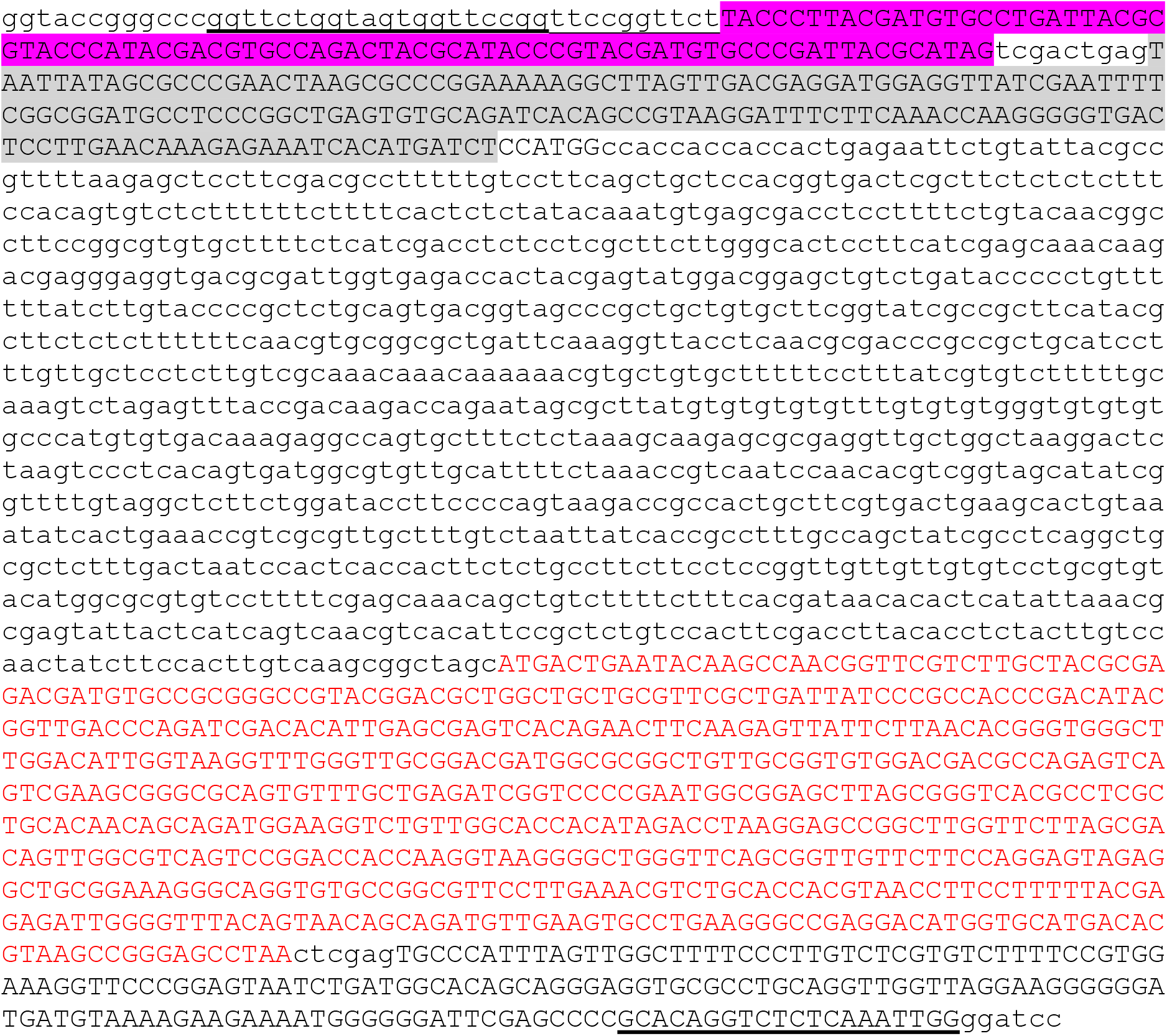

## Sequence of the *glmS*-tagging cassette in pMOTag-4-M9-puro-5

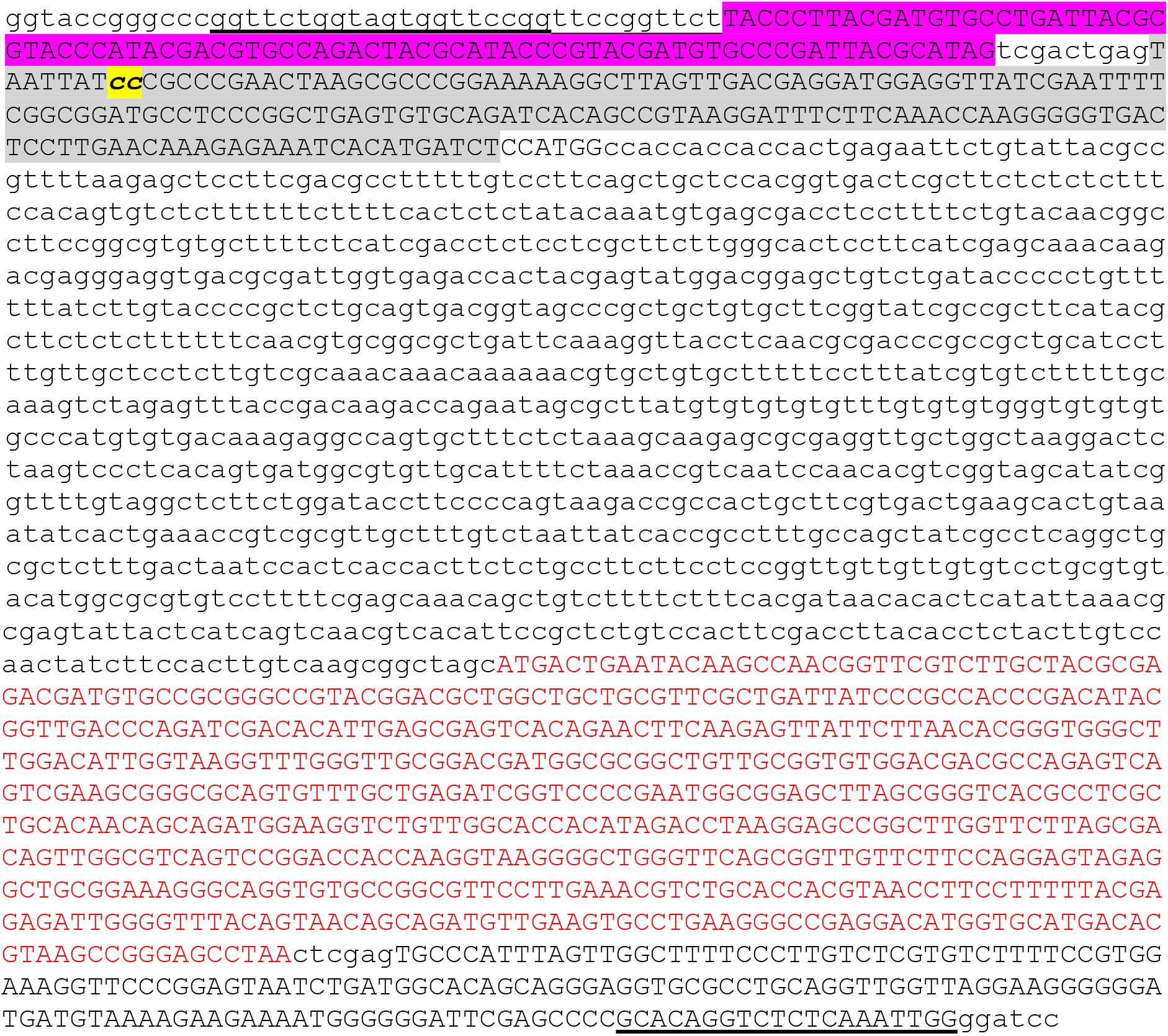

## Sequence of the *glmS*-tagging cassette in pMOTag-4-wt-blast-5

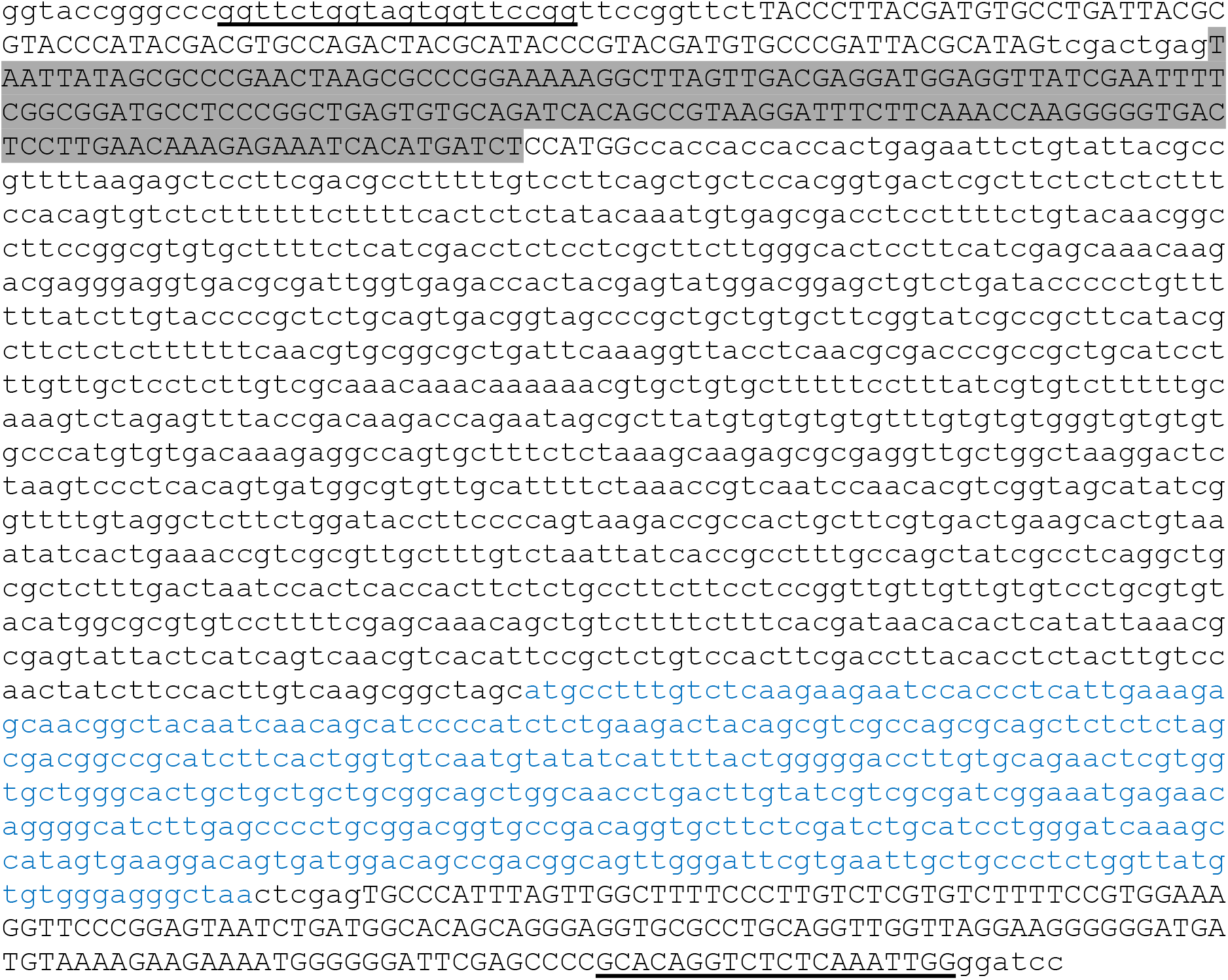

## Sequence of the *glmS*-tagging cassette in pMOTag-4-M9-blast-5

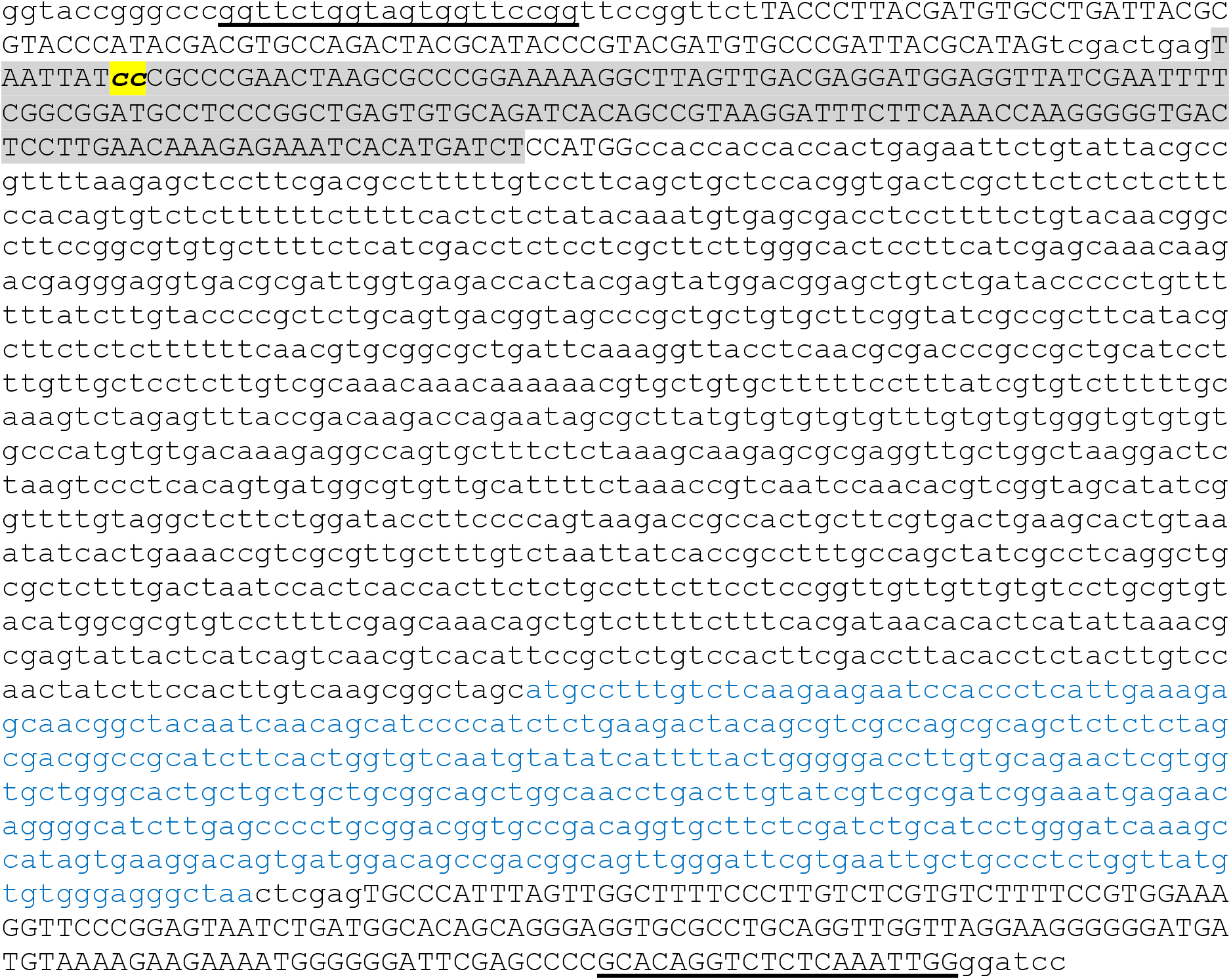

## References

1. World Health, O. Research priorities for Chagas disease, human African trypanosomiasis and leishmaniasis. World Health Organ Tech Rep Ser, v-xii, 1–100 (2012).

2. Global leishmaniasis update, 2006-2015: a turning point in leishmaniasis surveillance. Wkly Epidemiol Rec 92, 557–65 (2017).

3. Simpson, L. & Shaw, J. RNA editing and the mitochondrial cryptogenes of kinetoplastid protozoa. Cell 57, 355–66 (1989).

4. Ferguson, M.A. The structure, biosynthesis and functions of glycosylphosphatidylinositol anchors, and the contributions of trypanosome research. J Cell Sci 112 (Pt 17), 2799–809 (1999).

5. Li, F. et al. Structure of the core editing complex (L-complex) involved in uridine insertion/deletion RNA editing in trypanosomatid mitochondria. Proc Natl Acad Sci U S A 106, 12306–10 (2009).

6. Eckers, E., Cyrklaff, M., Simpson, L. & Deponte, M. Mitochondrial protein import pathways are functionally conserved among eukaryotes despite compositional diversity of the import machineries. Biol Chem 393, 513–24 (2012).

7. Breitling, R. et al. Non-pathogenic trypanosomatid protozoa as a platform for protein research and production. Protein Expr Purif 25, 209–18 (2002).

8. Kushnir, S., Gase, K., Breitling, R. & Alexandrov, K. Development of an inducible protein expression system based on the protozoan host Leishmania tarentolae. Protein Expr Purif 42, 37–46 (2005).

9. Lander, N., Li, Z.H., Niyogi, S. & Docampo, R. CRISPR/Cas9-Induced Disruption of Paraflagellar Rod Protein 1 and 2 Genes in Trypanosoma cruzi Reveals Their Role in Flagellar Attachment. mBio 6, e01012 (2015).

10. Zhang, W.W., Lypaczewski, P. & Matlashewski, G. Optimized CRISPR-Cas9 Genome Editing for Leishmania and Its Use To Target a Multigene Family, Induce Chromosomal Translocation, and Study DNA Break Repair Mechanisms. mSphere 2(2017).

11. Zhang, W.W. & Matlashewski, G. CRISPR-Cas9-Mediated Genome Editing in Leishmania donovani. MBio 6, e00861 (2015).

12. Beneke, T. et al. A CRISPR Cas9 high-throughput genome editing toolkit for kinetoplastids. R Soc Open Sci 4, 170095 (2017).

13. Beneke, T. et al. Genetic dissection of a Leishmania flagellar proteome demonstrates requirement for directional motility in sand fly infections. PLoS Pathog 15, e1007828 (2019).

14. Martel, D., Beneke, T., Gluenz, E., Spath, G.F. & Rachidi, N. Characterisation of Casein Kinase 1.1 in Leishmania donovani Using the CRISPR Cas9 Toolkit. Biomed Res Int 2017, 4635605 (2017).

15. Banaszynski, L.A., Chen, L.C., Maynard-Smith, L.A., Ooi, A.G. & Wandless, T.J. A rapid, reversible, and tunable method to regulate protein function in living cells using synthetic small molecules. Cell 126, 995–1004 (2006).

16. Deponte, M. GFP tagging sheds light on protein translocation: implications for key methods in cell biology. Cell Mol Life Sci 69, 1025–33 (2012).

17. Fire, A. et al. Potent and specific genetic interference by double-stranded RNA in Caenorhabditis elegans. Nature 391, 806–11 (1998).

18. Ngo, H., Tschudi, C., Gull, K. & Ullu, E. Double-stranded RNA induces mRNA degradation in Trypanosoma brucei. Proc Natl Acad Sci U S A 95, 14687–92 (1998).

19. Wilson, R.C. & Doudna, J.A. Molecular mechanisms of RNA interference. Annu Rev Biophys 42, 217–39 (2013).

20. Baum, J. et al. Molecular genetics and comparative genomics reveal RNAi is not functional in malaria parasites. Nucleic Acids Res 37, 3788–98 (2009).

21. Lye, L.F. et al. Retention and loss of RNA interference pathways in trypanosomatid protozoans. PLoS Pathog 6, e1001161 (2010).

22. Kolev, N.G., Tschudi, C. & Ullu, E. RNA interference in protozoan parasites: achievements and challenges. Eukaryot Cell 10, 1156–63 (2011).

23. Gossen, M. & Bujard, H. Tight control of gene expression in mammalian cells by tetracycline-responsive promoters. Proc Natl Acad Sci U S A 89, 5547–51 (1992).

24. Baron, U. & Bujard, H. Tet repressor-based system for regulated gene expression in eukaryotic cells: principles and advances. Methods Enzymol 327, 401–21 (2000).

25. Moullan, N. et al. Tetracyclines Disturb Mitochondrial Function across Eukaryotic Models: A Call for Caution in Biomedical Research. Cell Rep (2015).

26. Winkler, W.C., Nahvi, A., Roth, A., Collins, J.A. & Breaker, R.R. Control of gene expression by a natural metabolite-responsive ribozyme. Nature 428, 281–6 (2004).

27. McCarthy, T.J. et al. Ligand requirements for glmS ribozyme self-cleavage. Chem Biol 12, 1221–6 (2005).

28. Cochrane, J.C., Lipchock, S.V. & Strobel, S.A. Structural investigation of the GlmS ribozyme bound to Its catalytic cofactor. Chem Biol 14, 97–105 (2007).

29. Klein, D.J., Wilkinson, S.R., Been, M.D. & Ferre-D’Amare, A.R. Requirement of helix P2.2 and nucleotide G1 for positioning the cleavage site and cofactor of the glmS ribozyme. J Mol Biol 373, 178–89 (2007).

30. Prommana, P. et al. Inducible knockdown of Plasmodium gene expression using the glmS ribozyme. PLoS One 8, e73783 (2013).

31. Watson, P.Y. & Fedor, M.J. The glmS riboswitch integrates signals from activating and inhibitory metabolites in vivo. Nat Struct Mol Biol 18, 359–63 (2011).

32. Beneke, T. & Gluenz, E. LeishGEdit: A Method for Rapid Gene Knockout and Tagging Using CRISPR-Cas9. Methods Mol Biol 1971, 189–210 (2019).

33. Iovannisci, D.M., Goebel, D., Allen, K., Kaur, K. & Ullman, B. Genetic analysis of adenine metabolism in Leishmania donovani promastigotes. Evidence for diploidy at the adenine phosphoribosyltransferase locus. J Biol Chem 259, 14617–23 (1984).

34. Hwang, H.Y., Gilberts, T., Jardim, A., Shih, S. & Ullman, B. Creation of homozygous mutants of Leishmania donovani with single targeting constructs. J Biol Chem 271, 30840–6 (1996).

35. Specht, S. et al. A single-cysteine mutant and chimeras of essential Leishmania Erv can complement the loss of Erv1 but not of Mia40 in yeast. Redox Biol 15, 363–374 (2018).

36. Deponte, M. & Hell, K. Disulphide bond formation in the intermembrane space of mitochondria. J Biochem 146, 599–608 (2009).

37. Eckers, E. et al. Divergent molecular evolution of the mitochondrial sulfhydryl:cytochrome C oxidoreductase Erv in opisthokonts and parasitic protists. J Biol Chem 288, 2676–88 (2013).

38. Peikert, C.D. et al. Charting organellar importomes by quantitative mass spectrometry. Nat Commun 8, 15272 (2017).

39. Eckers, E. & Deponte, M. No need for labels: the autofluorescence of Leishmania tarentolae mitochondria and the necessity of negative controls. PLoS One 7, e47641 (2012).

40. Sollelis, L. et al. First efficient CRISPR-Cas9-mediated genome editing in Leishmania parasites. Cell Microbiol 17, 1405–12 (2015).

41. Basu, S. et al. Divergence of Erv1-associated mitochondrial import and export pathways in trypanosomes and anaerobic protists. Eukaryot Cell 12, 343–55 (2013).

42. Duncan, S.M., Jones, N.G. & Mottram, J.C. Recent advances in Leishmania reverse genetics: Manipulating a manipulative parasite. Mol Biochem Parasitol 216, 30–38 (2017).

43. Qi, L.S. et al. Repurposing CRISPR as an RNA-guided platform for sequence-specific control of gene expression. Cell 152, 1173–83 (2013).

44. Clayton, C.E. Life without transcriptional control? From fly to man and back again. EMBO J 21, 1881–8 (2002).

45. Daniels, J.P., Gull, K. & Wickstead, B. Cell biology of the trypanosome genome. Microbiol Mol Biol Rev 74, 552–69 (2010).

46. de Paiva, R.M. et al. Amastin Knockdown in Leishmania braziliensis Affects Parasite-Macrophage Interaction and Results in Impaired Viability of Intracellular Amastigotes. PLoS Pathog 11, e1005296 (2015).

47. Robinson, K.A. & Beverley, S.M. Improvements in transfection efficiency and tests of RNA interference (RNAi) approaches in the protozoan parasite Leishmania. Mol Biochem Parasitol 128, 217–28 (2003).

48. DaRocha, W.D., Otsu, K., Teixeira, S.M. & Donelson, J.E. Tests of cytoplasmic RNA interference (RNAi) and construction of a tetracycline-inducible T7 promoter system in Trypanosoma cruzi. Mol Biochem Parasitol 133, 175–86 (2004).

49. Madeira da Silva, L., Owens, K.L., Murta, S.M. & Beverley, S.M. Regulated expression of the Leishmania major surface virulence factor lipophosphoglycan using conditionally destabilized fusion proteins. Proc Natl Acad Sci U S A 106, 7583–8 (2009).

50. Wheeler, R.J., Gluenz, E. & Gull, K. Basal body multipotency and axonemal remodelling are two pathways to a 9+0 flagellum. Nat Commun 6, 8964 (2015).

51. Damerow, S. et al. Depletion of UDP-Glucose and UDP-Galactose Using a Degron System Leads to Growth Cessation of Leishmania major. PLoS Negl Trop Dis 9, e0004205 (2015).

52. Podesvova, L., Huang, H. & Yurchenko, V. Inducible protein stabilization system in Leishmania mexicana. Mol Biochem Parasitol 214, 62–64 (2017).

53. Yan, S., Myler, P.J. & Stuart, K. Tetracycline regulated gene expression in Leishmania donovani. Mol Biochem Parasitol 112, 61–9 (2001).

54. Kraeva, N., Ishemgulova, A., Lukes, J. & Yurchenko, V. Tetracycline-inducible gene expression system in Leishmania mexicana. Mol Biochem Parasitol 198, 11–3 (2014).

55. Counihan, N.A. et al. Plasmodium falciparum parasites deploy RhopH2 into the host erythrocyte to obtain nutrients, grow and replicate. Elife 6 (2017).

56. Bridgford, J.L. et al. Artemisinin kills malaria parasites by damaging proteins and inhibiting the proteasome. Nat Commun 9, 3801 (2018).

57. Marapana, D.S. et al. Plasmepsin V cleaves malaria effector proteins in a distinct endoplasmic reticulum translocation interactome for export to the erythrocyte. Nat Microbiol 3, 1010–1022 (2018).

58. Hallee, S. et al. Identification of a Golgi apparatus protein complex important for the asexual erythrocytic cycle of the malaria parasite Plasmodium falciparum. Cell Microbiol 20, e12843 (2018).

59. Cruz-Bustos, T., Ramakrishnan, S., Cordeiro, C.D., Ahmed, M.A. & Docampo, R. A Riboswitch-based Inducible Gene Expression System for Trypanosoma brucei. J Eukaryot Microbiol 65, 412–421 (2018).

60. Lander, N., Cruz-Bustos, T. & Docampo, R. A CRISPR/Cas9-riboswitch-Based Method for Downregulation of Gene Expression in Trypanosoma cruzi. Front Cell Infect Microbiol 10, 68 (2020).

61. LeBowitz, J.H., Coburn, C.M., McMahon-Pratt, D. & Beverley, S.M. Development of a stable Leishmania expression vector and application to the study of parasite surface antigen genes. Proc Natl Acad Sci U S A 87, 9736–40 (1990).

62. Simpson, L., Frech, G.C. & Maslov, D.A. RNA editing in trypanosomatid mitochondria. Methods Enzymol 264, 99–121 (1996).

63. Schumann Burkard, G., Jutzi, P. & Roditi, I. Genome-wide RNAi screens in bloodstream form trypanosomes identify drug transporters. Mol Biochem Parasitol 175, 91–4 (2011).

